# Impact of histone deacetylase inhibition and arimoclomol on heat shock protein expression and disease biomarkers in primary culture models of familial ALS

**DOI:** 10.1101/2023.12.13.571549

**Authors:** Mario Fernández Comaduran, Sandra Minotti, Suleima Jacob-Tomas, Javeria Rizwan, Nancy Larochelle, Richard Robitaille, Chantelle F. Sephton, M. Vera, Josephine N. Nalbantoglu, Heather D. Durham

**Author notes:** **Corresponding author:** Heather D. Durham, Rm 649, Montreal Neurological Institute, 3801 University St. Montreal, QC, H3A 2B4 Canada.

## Abstract

Protein misfolding and mislocalization are common themes in neurodegenerative disorders, including the motor neuron disease, amyotrophic lateral sclerosis (ALS). Maintaining proteostasis is a crosscutting therapeutic target, including upregulation of heat shock proteins (HSP) to increase chaperoning capacity. Motor neurons have a high threshold for upregulating stress inducible HSPA1A, but constitutively express high levels of HSPA8. This study compared expression of these HSPs in cultured motor neurons expressing three variants linked to familial ALS: TDP-43^G348C^, FUS^R521G^ or SOD1^G93A^. All variants were poor inducers of *Hspa1a,* and reduced levels of *Hspa8* mRNA and protein, indicating multiple compromises in chaperoning capacity. To promote HSP expression, cultures were treated with the putative HSP co-inducer, arimoclomol, class I histone deacetylase (HDAC) inhibitors to promote active chromatin for transcription, and the combination. Treatments had variable, often different effects on expression of *Hspa1a* and *Hspa8*, depending on the ALS variant expressed, mRNA distribution (somata and dendrites), and biomarker of toxicity measured (histone acetylation, maintaining nuclear TDP-43 and the nBAF chromatin remodeling complex component Brg1, mitochondrial transport, FUS aggregation). Overall, HDAC inhibition alone was effective on more measures than arimoclomol. In the TDP-43 model, arimoclomol failed to induce HSPA1A or preserve *Hspa8* mRNA, despite preserving nuclear TDP-43 and Brg1, indicating neuroprotective properties other than HSP induction. The data speak to the complexity of drug mechanisms against multiple biomarkers of ALS pathogenesis, as well as to the importance of HSPA8 for neuronal proteostasis in both somata and dendrites.

## Introduction

Heat shock proteins (HSPs) act as protein chaperones to assist protein folding and to maintain proteome stability under physiological and stress conditions. They recognize misfolded proteins or unstable intermediates, interact with them to prevent inappropriate interactions, and refold or target them for degradation. A general biological principle is that cells regulate expression of heat shock genes according to demand and upregulate them under stress, the so-called heat shock response. This response can be compromised by aging and disease. Accumulation of misfolded proteins and protein aggregation are a common theme in neurodegenerative disorders, including amyotrophic lateral sclerosis (ALS). A logical therapeutic strategy is to improve the function of pathways involved in maintaining a healthy proteome, including upregulation of the chaperone network; however, this approach has been complicated by the high threshold for activating transcription of stress-inducible heat shock genes in neurons, including motor neurons, and loss of efficacy with disease progression consequent to disease mechanisms, including changes in chromatin structure [1–3].

The drug arimoclomol has been investigated as a heat shock protein co-inducer; *i.e*., to magnify an existing stress response by prolonging interaction of the major heat shock transcription factor, HSF1, with heat shock elements (HSE) in heat shock gene promoters [4]. Arimoclomol showed some efficacy in a mouse model of familial ALS (fALS) due to mutation in the superoxide dismutase I (*SOD1*) gene (ALS1) [5] and promise in ALS patients with *SOD1* mutation [6], but it failed to meet primary endpoints in a [clinical trial](https://www.clinicaltrials.gov/study/NCT03836716?cond=als&term=ORARIALS-01&rank=1&tab=results) in sporadic ALS patients.

We have been investigating approaches to facilitate upregulation of heat shock genes in motor neurons and recently reported that histone deacetylase (HDAC) inhibitors enabled the heat shock response in motor neurons of dissociated murine spinal cord cultures, improving the efficacy of HSP-inducing drugs in response to proteotoxic stress induced by expression of the SOD1^G93A^ variant [7]. Consistent with our previous report [8], the stress-inducible HSP70 isoform, HSPA1A, was induced in only about 10% of motor neurons expressing SOD1^G93A^. Although arimoclomol was a poor coinducer, efficacy was enhanced by combined treatment with a histone deacetylase (HDAC) inhibitor, particularly through inhibition of class I HDACs [7].

HDAC inhibition strongly accentuated the induction of HSPA1A expression by HSP90 inhibitors [7]. HSF1 is maintained in an inactive state as part of HSP90 complexes, like many other transcription factors. Under stress or when treated with HSP90 inhibitors that disrupt the complex, HSF1 is released to engage with heat shock elements (HSE) of heat shock genes [9]. HDAC inhibition could increase chromatin accessibility by maintaining histone residues in the acetylated state associated with the ‘openness’ of chromatin and accessibility of transcription factors.

HSF1-HSE interaction is also influenced by SWI/SNF chromatin remodeling complexes in yeast [10]. The mammalian ortholog, the neuronal Brm/Brg-associated factor (nBAF) chromatin remodeling complex, drives neuronal differentiation in development and maintains dendritic architecture in mature neurons [11]. Dendritic attrition and loss of nBAF complexes in motor neurons are features of ALS, implicating chromatin remodeling pathways in ALS [12,13].

ALS6 is caused by dominantly inherited mutations in the gene encoding fused in sarcoma (FUS) [14]. When expressed in cultured spinal motor neurons, FUS variants caused features of the disease including loss of nuclear FUS and accumulation and aggregation in the cytoplasm, as well as loss of dendritic branching associated with loss of nBAF chromatin remodeling complexes [13,15]. In motor neurons expressing FUS^R521G/H^, HDAC inhibition preserved nuclear FUS [7]. Arimoclomol was as efficient as HDAC inhibition in preserving nuclear FUS, despite lack of direct class I HDAC inhibitory activity, and combination treatment with HDAC inhibitor and arimoclomol was even more effective. Neither expression of the FUS variant nor treatment with HDAC inhibitors and/or arimoclomol induced expression of HSPA1A, indicating that the heat shock response was even further suppressed in motor neurons expressing FUS variants in comparison to SOD1^G93A^ [7].

Those data encouraged us to pursue studies to test arimoclomol in combination with a class I HDAC inhibitor in ALS models. An important limitation with most commercially available HDAC inhibitors is target engagement in the CNS. RGFP963 was obtained from BioMarin as a class I HDAC inhibitor with CNS biodistribution. In the present study, RGFP963 was included, alone and in combination with arimoclomol, for induction of heat shock proteins in cultured motor neurons expressing ALS-linked variants and for impact of the treatments on multiple biomarkers of toxicity.

We focused experiments on TDP-43 variants linked to fALS as well as pursuing experiments with SOD1 and FUS variants. ALS10 is caused by dominantly inherited mutations in the *TARDBP* gene encoding TAR DNA Binding Protein 43 kDa (TDP-43), a DNA/RNA-binding protein normally concentrated in the nucleus but shuttling between the nucleus and cytoplasm [16]. Depletion of TDP-43 from the nucleus and accumulation in cytoplasmic inclusions can be found in autopsy specimens in the majority of ALS cases and in frontotemporal dementia (FTD) [17–19]. Although *TARDBP* mutations account for a small percentage of ALS cases, the general presence of this pathology in other ALS types indicates an important role of the protein in stressed neurons and in ALS pathogenesis [17,20]. Our previous study established a culture model of ALS10, showing depletion of nuclear TDP-43, dendritic attrition and loss of nBAF chromatin remodeling complexes in motor neurons expressing TDP-43^G348C^, similar to neurons expressing FUS variant.

Here we report neuroprotective properties of both HDAC inhibitors and arimoclomol in motor neurons in all three fALS culture models that were not linked to upregulation of HSPA1A. This stress-inducible HSP is typically used as a marker of the heat shock response, due to low or absent expression under homeostatic conditions and upregulation during stress. However, constitutively expressed HSPs are the work horse in maintaining protein quality under physiological conditions. Motor neurons express high levels of HSC/HSP70s and I.R. Brown *et al*. proposed that neurons would rely on these isoforms when under stress, in the absence of HSPA1A [21]. Therefore, we investigated the expression of the heat shock cognate protein 70 (HSC70; HSPA8). *Hspa8* mRNA levels declined in somata and dendrites of motor neurons expressing ALS variants. RGFP963 prevented this loss in the SOD1 model, but not in motor neurons expressing the FUS variant; only the combination of RGFP963 and arimoclomol preserved somal, but not dendritic, *Hspa8*. These data point to differences in how ALS variants affect chaperone expression and to neuroprotective properties of arimoclomol through other mechanisms.

## Materials and methods

### fALS culture models

Preparation and maintenance of dissociated spinal cord-DRG cultures from E13 CD1 mouse embryos (Charles River Laboratories, St. Constant, QC, Canada) were as previously described [22]. Briefly, ∼450,000 Cells were seeded onto 18 mm glass coverslips (Thermo Fisher Inc., St. Laurent, QC, Canada) previously coated with poly-D-lysine (Millipore Sigma, Oakville, ON, Canada) and Matrigel® basement membrane matrix (Millipore Sigma) in 12-well culture dishes (Greiner Bio-One, Frickenhausen, Germany). Cultures were maintained at 37⁰C and 5% CO_2_ in the modified N3 growth medium described in Roy et al. (1998) plus 1.3% horse serum (Invitrogen Canada Inc., Burlington, ON, Canada), 5 ng/mL 2.5S nerve growth factor (Millipore) and 0.5X B-27 Supplement (Life Technologies, Carlsbad, CA, USA). At confluency, cultures were treated with 1.4 μg/mL cytosine-β-D-arabinofuranoside (Calbiochem) for three days to suppress division of non-neuronal cells. Cultures were matured for at least three weeks for development of motor neurons (Fig. 1A), identified as described in Roy et al. (1998).

**Figure 1.**
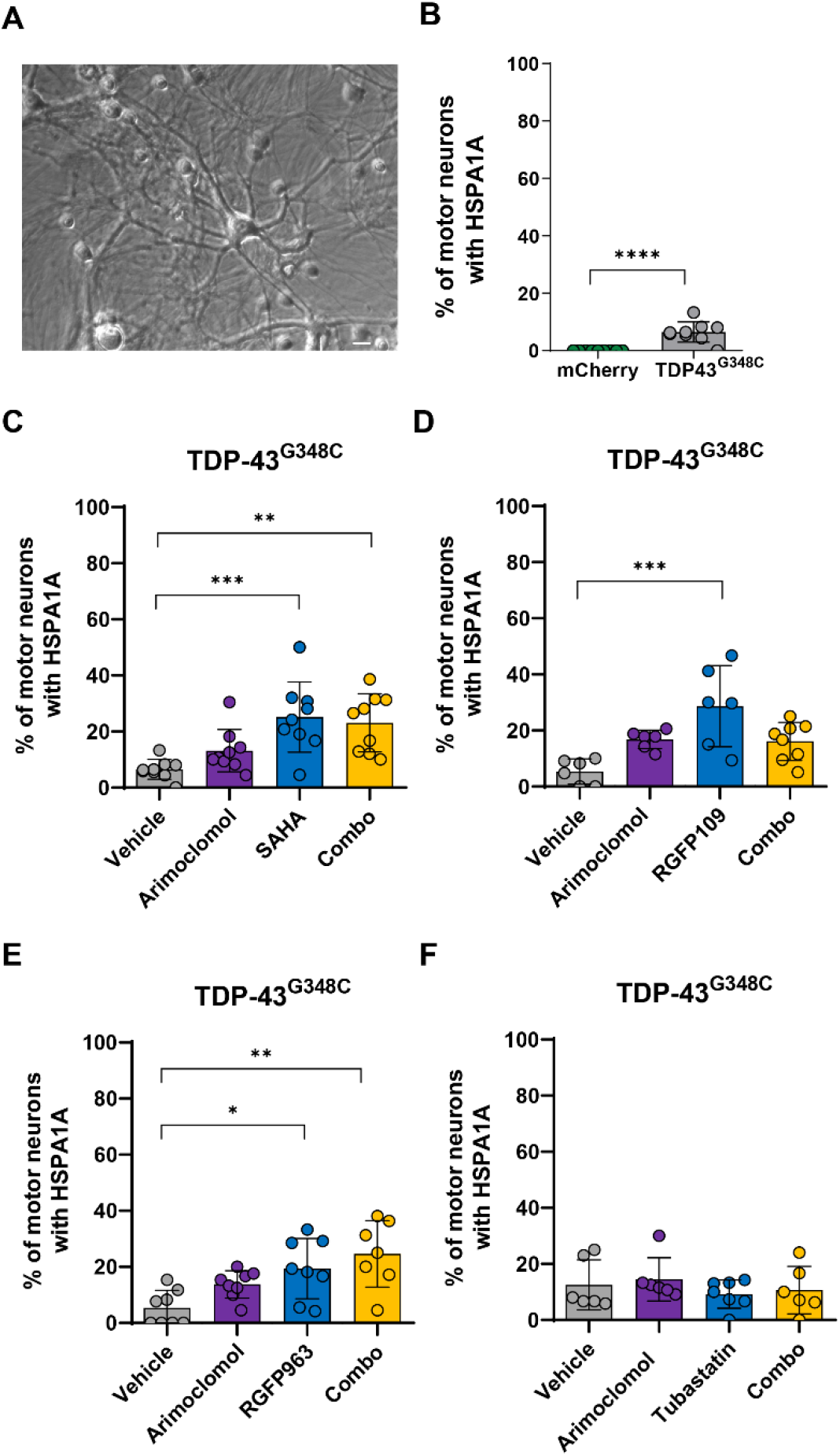
Effects of HDAC inhibitors, arimoclomol and combined treatments on HSPA1A in motor neurons expressing TDP-43^G348C^. **A** Phase contrast image of spinal cord – DRG culture. Arrows point to motor neurons. **B** Small but significant increase in the percentage of motor neurons with HSPA1A immunoreactivity three days following microinjection of plasmid encoding TDP-43^G348C^, compared to absence of labeling in neurons injected with mCherry (control). **C-F** Percentage of HSPA1A immunopositive motor neurons after three days of treatment with vehicle, the HDAC inhibitors **C** SAHA, **D** RGFP109, **E** RGFP963 or **F** Tubastatin A, or with arimoclomol alone or in combination with the respective HDAC inhibitor. HDAC inhibitors with class I activity (**C-E**) significantly increased the percentage of neurons expressing HSPA1A. Arimoclomol and the HDAC6 inhibitor Tubastatin A were ineffective. Data presented as mean ± S.D., n = 9 cultures. Statistical significance was evaluated by one-way ANOVA followed by Bonferroni post hoc analysis. *p<0.05, **p<0.01, ***p<0.001, ****p<0.0001. Scale bar = 20μm.

fALS culture models were established by intranuclear microinjection of plasmids encoding ALS variants and/or cell markers into motor neuronal nuclei as previously described [23,24]. Plasmids used were human *TARDBP* (WT or G348C) with N-terminus Flag and C-terminus Myc tags in pCS2+ microinjected at 20 ng/µL [25], N-terminal Flag-tagged human FUS (WT or R521G) in pcDNA3 at 20 ng/μL [15], SOD1 (WT or G93A) in pCEP4 at 200 ng/μL or pcDNA3 at 27 ng/μL [23], mCherry-C1 plasmid (Clontech, Mountain View, CA, USA) at 1 ng/μl, eGFP plasmid (Clontech) at 5 ng/μL and pOCTmCherry at 1 μg/mL (Tradewell et al. 2011).

### Immunocytochemistry

Cultures on coverslips were fixed in 4% paraformaldehyde (PFA) in PBS for 10 min; permeabilized using 0.5% NP-40 for 5 min; resubmerged in 4% PFA in PBS for 2 min; blocked in 5% horse serum in PBS for 30 min at R.T. or overnight at 4°C; incubated with primary antibody for 1 hr; washed three X 5 min in PBS; incubated with secondary antibody for 45 min; washed three X 5 min in PBS, and mounted on microscope slides in Epredia™ Immu-Mount™ (Thermo Fisher Scientific Inc.).

*Primary antibodies* were: rabbit anti-TDP-43 (EPR18554 1:500 Abcam, Cambridge, UK), rabbit anti-FUS (11570-1-AP 1:300, Proteintech, Rosemont, IL, USA); mouse antibody specific for human SOD1 (SD-G6 1:100, Millipore Sigma), Mouse anti-human HSP70 specific for stress-inducible HSPA1A (SMC-100B 1:100, Stressmarq Biosciences Inc., Victoria, BC, Canada), mouse anti-HSPA8/HSC71/Hsc70 (13D3) antibody (NB120-2788 1:300, Novus Biologicals, Colorado, USA), mouse anti-FLAG M2 (F1804 1:400, Millipore Sigma), rabbit anti-acetylated H3K9/K14 (9677 1:400, Cell Signaling, Danvers, MA, USA), rabbit anti-Brg1 (21634-1-AP 1:600, Proteintech), rabbit anti-MAP2 (AB5622 1:500, Millipore Sigma), chicken anti-GFP (GFP1010 1:500 AVES, CA, USA).

*Secondary antibodies* for immunofluorescence (Jackson Immunoresearch, Cedarlane, Burlington, ON, Canada, unless stated otherwise): Alexa Fluor 488-conjugated donkey anti-mouse IgG (1:300), Cy5-, Cy3- and Cy2-conjugated donkey anti-rabbit IgG (1:300), Cy3-donkey anti-mouse IgG (1:300), Cy2-donkey anti-mouse IgG (Rockland, PA, USA; 610-711-124; 1:300), Alexa Fluor 750-goat anti-rabbit (Invitrogen; 1:1000), Alexa Fluor 488-goat anti-chicken (Invitrogen; 1:1000), and Alexa 750-goat anti mouse (Invitrogen; 1:1000).

*Imaging* was carried out using a Zeiss Observer Z1 microscope (Carl Zeiss Microscopy) equipped with a Hamamatsu ORCA-ER cooled CD camera (Hamamatsu, Japan). Images were acquired and analyzed with Zeiss Axiovision software.

### Drug treatments

Cultures were treated post microinjection and analyzed after three days, unless otherwise stipulated. Drugs used were: Arimoclomol (Medkoo Biosciences, Morrisville, NC, USA) dissolved in water to prepare a stock solution of 2 mM; HDAC inhibitors: the pan HDAC inhibitor suberoylanilide hydroxamic acid [SAHA] (Cedarlane), HDAC 1/3 inhibitors RGFP963 (BioMarin Pharmaceutical Inc., San Raphael, CA, USA) and RGFP109 (Selleck Chemicals LLC, Houston, TX, USA) dissolved in DMSO for stock solutions of 2mM, and the HDAC6 inhibitor Tubastatin A (Cayman Chemical, Ann Arbor, MI, USA) dissolved in DMSO for a stock solution of 6 mM. Drugs were diluted to working concentrations in modified N3 culture medium with gentamycin, with equivalent concentration of DMSO as control.

### Quantifying HSP mRNA expression using single-molecule fluorescence *in situ* hybridization (smFISH)

Motor neurons were microinjected with plasmids encoding ALS variant and eGFP to identify injected cells. On day one, two or three post microinjection, a combined smFISH with immunofluorescence protocol was followed as described in detail by Jacob-Tomas et al. [26], adapted from Eliscovich et al. [27]. The probes used for mus musculus *Hspa1a* or *Hspa8* mRNA (Stellaris LGC Biosearch Technologies, CA, USA) are a mixture of up to 48 lyophilized DNA oligonucleotides, each of them being between 18 and 22 nucleotides in length [26]. The sequence of probes labeled with Quasar 570 or Quasar 670 are presented in Supplementary Fig. 1.

After prehybridization and blocking, cells were incubated with smFISH *Hspa1a* and *Hspa8* probes at 125 nM and primary antibodies against GFP, MAP2 to identify dendrites and SOD1, TDP-43 or FUS in hybridization buffer, incubated at 37°C for 3 hr, washed 2X using warmed prehybridization buffer and then incubated with secondary antibodies in prehybridization buffer for 1 hr at 37°C in the dark. After washing, coverslips were air dried and mounted in ProLong Gold Antifade Reagent (Thermo Fisher Scientific Inc.). The smFISH and epifluorescent signals were detected using a customized inverted Nikon Ti-2 wide-field microscope using a 60X 1.4NA oil immersion objective (Nikon), Spectra X LED light engine with a Cy7+ bandpass filter (Lumencor, Beaverton, OR), and Orca-Fusion sCMOS camera (Hamamatsu). Automated Nikon Elements software was used to acquire a stack of 41 optical planes (at 0.2 μm steps) consecutively in 5 channels: *Hspa1a* mRNA – 555nm; *Hspa8* mRNA – 640nm; TDP-43, FUS or SOD1 – 750nm; GFP or MAP2 – 470 nm; DAPI 395nm), with x-y pixel size of 107.5nm, and z-step size of 200nm.

The subcellular distribution of epifluorescent spots from the mRNA probe, representing single mRNAs, was analyzed using a specialized protocol developed by the Vera lab, employing a segmented computational approach using Python and FISH-Quant [26]. See Wefers *et al*. for detailed methods [28]. Separate analysis of somal cytoplasm and dendritic projections was carried out using the FISH-Quant polygon tool to outline regions of interest. Foci of probe labeling in the nucleus representing transcription sites also were quantified.

### Quantifying intensity of HSPA8 immunolabeling

Three days following microinjection with plasmid encoding ALS variants, cultures were double labeled with mouse anti-HSPA8 antibody and antibody to TDP-43, SOD1, or FUS, using Alexa Fluor 488 anti-rabbit and Cy3 anti-mouse secondary antibodies. Non-injected motor neurons served as controls. Images were acquired as described for smFISH, converted to 16-bit format using ImageJ and then annotated to identify the soma. The resulting annotated images were processed using Python to generate print files that served as outlines. These outlines were further analyzed for pixel intensity using a Python code developed for this purpose.

### Mitochondrial transport

Mitochondrial transport was assessed by live imaging three days following microinjection of plasmids encoding ALS variant and pOCT-mCherry (the ornithine carbamoyltransferase signal sequence targeting mCherry to mitochondria) using the Zeiss Observer Z1 microscope, Hamamatsu ORCA-ER cooled CD and Axiovision software described above. Time series acquisitions of mCherry epifluorescence in initial axonal segments of motor neurons were configured to record frames at 3 sec intervals over a 7 min interval. Fiji ImageJ (NIH) was used to generate kymographs of mitochondrial movement using the KymoResliceWide plug-in. Kymographs were analyzed using the KymoButler plug-in, which automatically tracks dynamic processes in kymographs [29], designed for Mathematica (Wolfram Research, Inc.). After visual examination for artifacts, mitochondrial velocities determined by the KymoButler plug-in were extracted and subjected to statistical analysis.

### Statistical analysis

Statistical analyses were performed in Prism 8 (GraphPad Software Inc.); *i.e.,* unpaired Student’s t-tests to compare two data sets and one-way ANOVA followed by Bonferroni’s or Dunnett’s multiple comparison tests to compare more than two data sets. Significance was established at p < 0.05.

## Results

### Effect of fALS variants, HDAC inhibition and arimoclomol on the heat shock response in cultured motor neurons

#### HSPA1A in motor neurons expressing fALS-linked variants

Experimental models of ALS in which fALS-linked variants are expressed in motor neurons have the advantage of examining these variants and candidate therapies in the context of the unique physiology that increases motor neuron vulnerability to disease. We previously reported that SOD1^G93A^, causing fALS1, demonstrated limited capacity to induce expression of the stress-inducible HSP70 isoform, HSPA1A, in cultured motor neurons [7,8]. The percentage of motor neurons expressing HSPA1A three days following microinjection of expression plasmid was significantly increased when cultures were exposed to arimoclomol, to the pan HDAC inhibitor SAHA or the class I HDAC inhibitor RGFP109, but not the HDAC6 inhibitor Tubastatin A [7]. Although the combination of arimoclomol and SAHA was significantly more effective than either drug alone, most neurons remained resistant. No expression of HSPA1A was detected in motor neurons expressing the fALS6-linked variant FUS^R521G^ even with drug treatments.

#### Effect of TDP-43^G348C^ on HSPA1A expression and the consequences of drug treatments

TDP-43^G348C^ also is a poor inducer of the heat shock response. HSPA1A was detected by immunolabeling in a mean of only 6.5% of cultured motor neurons expressing this variant 3 days following intranuclear microinjection of expression plasmid, in contrast to absence of labeling in neurons expressing mCherry (control) (Fig. 1B). The pan HDAC inhibitor, SAHA, (Fig. 1C) and class I HDAC inhibitors, RGFP109 (Fig. 1D) and RGFP963 (Fig. 1E), significantly increased the percentage of motor neurons with HSPA1A immunolabeling, whereas the HDAC6 inhibitor Tubastatin A had no effect (Fig. 1F). Qualitatively, those results resemble those obtained in neurons expressing SOD1^G93A^, except that arimoclomol was not effective in neurons expressing TDP-43^G348C^, nor did it improve the efficacy of HDAC inhibitors when used in combination.

#### *Hspa1a* mRNA levels and cellular distribution in motor neurons expressing fALS variants

Expression of *Hspa1a* mRNA, measured by smFISH on day three post microinjection, was consistent with the low protein level. eGFP, co-expressed with ALS variant for visual identification of injected neurons as well as serving as procedural control, did not induce expression of *Hspa1a* mRNA. Fig. 2A illustrates a motor neuron labeled with probe specific for the constitutively expressed *Hspa8* for comparison to the *Hspa1a* probe (Fig. 2B,C,D). Few neurons microinjected with plasmid encoding TDP-43^G348C^ (Fig. 2B), FUS^R521G^ (Fig. 2C) or SOD1^G93A^ (Fig. 2D) expressed *Hspa1a* mRNA. Treatment with arimoclomol or RGFP963 increased the percentage of motor neurons expressing this mRNA only in the SOD1 model, but as with protein measurement, most neurons remained refractory.

**Figure 2.**
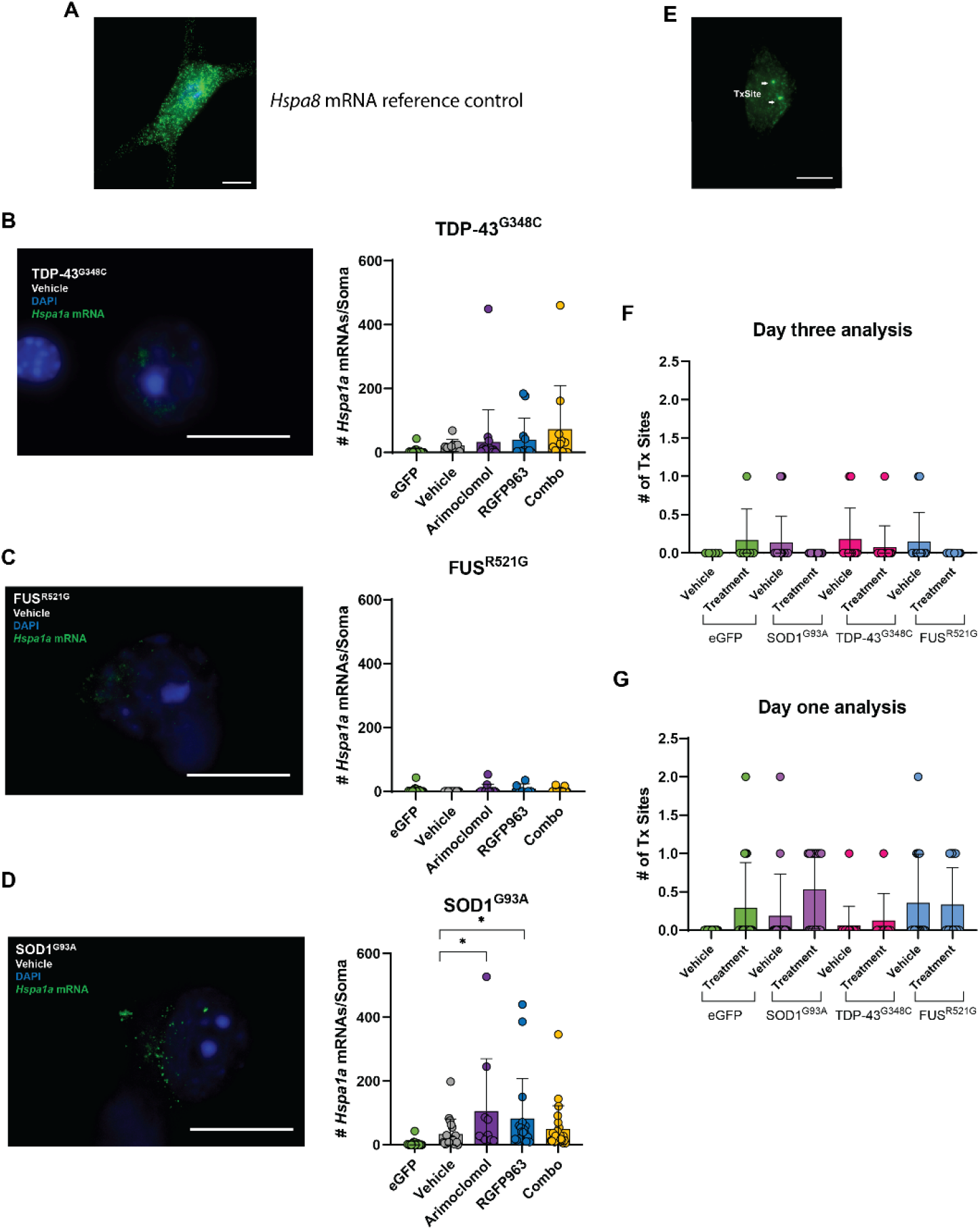
*Hspa1a* mRNA in ALS culture models using single molecule fluorescence *in situ* hybridization (smFISH). Cultures containing motor neurons expressing TDP-43^G348C^, FUS^R521G^ or SOD1^G93A^ were treated with vehicle (DMSO), 4 µM arimoclomol, 1µM RGFP963 or the combination of arimoclomol and RGFP963 (Combo). smFISH was conducted on day three. **A** The highly expressed *Hspa8* mRNA is presented as a reference control for comparison to the minimal labeling of *Hspa1a* mRNA in **B,C,D**. **B** No significant impact of TDP-43^G348C^ or drug treatments on the number of *Hspa1a* mRNA spots. **C** Scarcity of *Hspa1a* mRNA spots in neurons expressing FUS^R521G^; no significant effect of drug treatments. **D** Low numbers of *Hspa1a* mRNA spots in neurons expressing SOD1^G93A^; a small but significant increase in a subset of neurons by arimoclomol and RGFP963 treatments. Data are presented as mean ± S.D. n = 9-24 neurons per group. **E-G** Number of transcription sites by smFISH. **E** Example of H*spa1a* transcription site labeling. **F,G** Effect of ALS variants and combination drug treatment on the number of *Hspa1a* transcription sites on **F** day one and **G** day three following microinjection of expression vectors. n = 6-30 neurons per group. Statistical significance was evaluated through one-way ANOVA followed by Bonferroni post hoc analysis. *p<0.05. Scale bar = 15μm.

#### *Hspa1a* transcription sites

The number of transcription sites, illustrated in Fig. 2E, were quantified on day one (Fig. 2F) and on day three (Fig. 2G) post microinjection. Few motor neurons expressing eGFP, SOD1^G93A^, TDP-43^G348C^ or FUS^R521G^ had visible transcription sites, there being no significant difference between groups; however, on day one, a few motor neurons had up to two transcription sites, whereas on day three no motor neurons had more than one (8C), consistent with a weak but unsustained transcriptional activation in a subset of neurons. The number of motor neurons with *Hspa1a* transcription sites was not increased by treatment with arimoclomol in combination with RGFP963. These results point to the major cause of poor induction of HSPA1A by proteotoxic stress in motor neurons being at the level of transcription.

The data reinforce the differences in impact ALS-linked variants have on the heat shock response and have implications for therapy with HSP co-inducers, which depend on an initial activation of the heat shock response to be effective.

#### Impact of ALS variants on the constitutively expressed HSC, HSPA8

The high level of constitutively expressed HSP/HSC isoforms in motor neurons is likely to serve as a defense mechanism against proteotoxic stress [21]. Given the poor induction of HSPA1A, we turned our attention to the effect of ALS variants and of arimoclomol and RGFP963 on the major HSC isoform, HSPA8.

*Hspa8* mRNA spots, representing individual mRNA molecules, were quantified in both the soma and dendrites of motor neurons, given the importance of transport of HSP-encoding mRNAs to dendrites and dendritic spines to maintaining local protein quality control [30] (Fig. 3). *Hspa8* mRNA was highly expressed in somata of control motor neurons (injected neurons expressing eGFP) (mean of 1370 mRNA spots); however, a significant reduction occurred in motor neurons expressing SOD1^G93A^ by day three (mean of 580 *Hspa8* mRNA spots). Similarly, motor neurons expressing TDP-43^G348C^ displayed a reduced mean count of 590 mRNA spots, and even lower expression was measured in those expressing FUS^R521G^ (mean of 490 mRNA spots) (Fig. 3B). A significant reduction in dendritic mRNAs also occurred in motor neurons expressing ALS variants, from a mean count of 160 *Hspa8* mRNA spots in control, to 60 mRNAs in neurons expressing SOD1^G93A^, 70 mRNAs in neurons expressing TDP-43^G348C^ and 50 mRNAs in neurons expressing FUS^R521G^ (Fig. 3C). Wild type SOD1, TDP-43 and FUS were also investigated (Fig. 3D,E). Only TDP-43 overexpression reduced *Hspa8* mRNA, consistent with many other studies showing toxicity of its upregulation.

**Figure 3.**
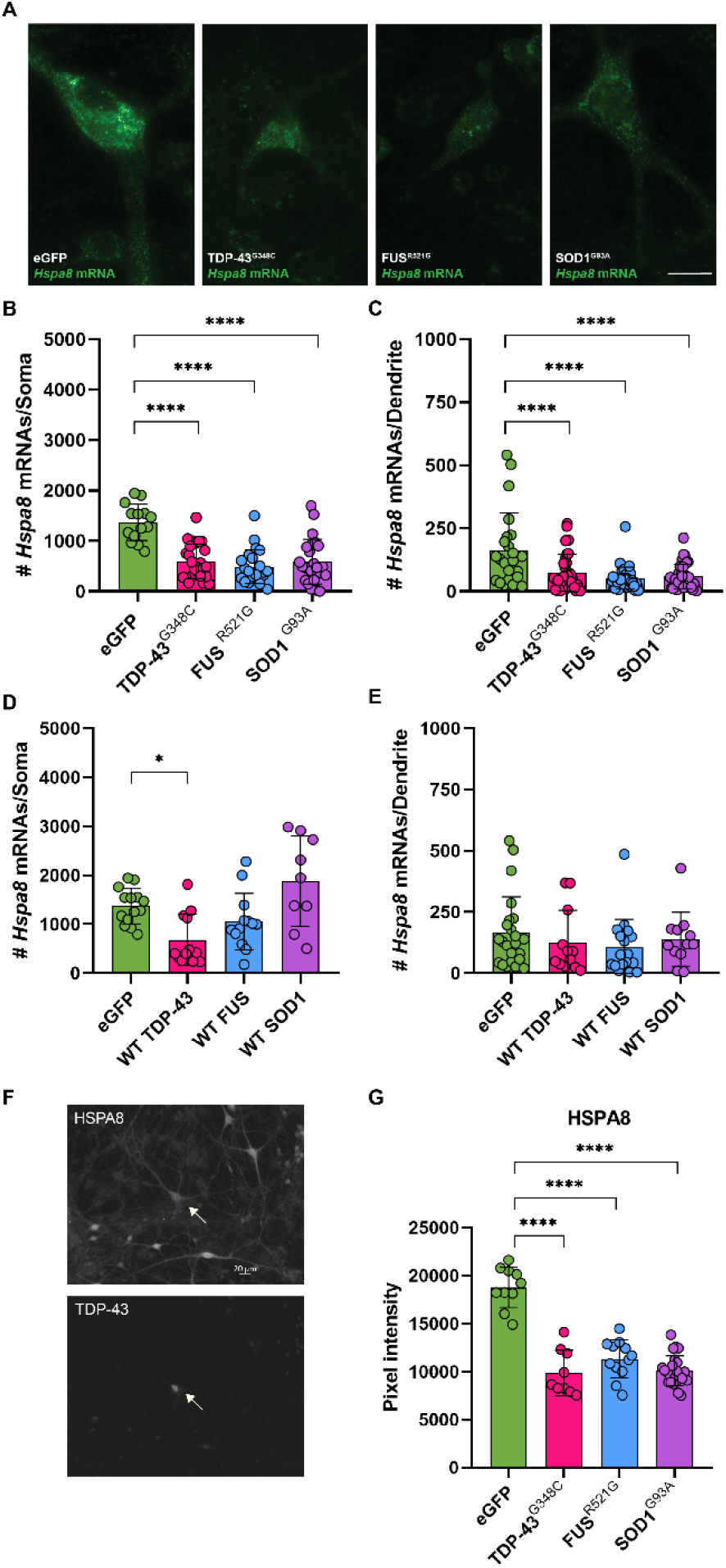
**A-E** ALS variants suppress *Hspa8* mRNA levels in motor neurons. *Hspa8* mRNA was quantified by smFISH three days after microinjection of TDP-43^G348C^, FUS^R521G^ or SOD1^G93A^ plus eGFP vectors. Microinjected motor neurons were identified by double-immunolabeling with antibodies to eGFP and TDP-43, FUS or SOD1. **A** Examples of *Hspa8* smFISH. Significant decrease in RNA spot counts in **B** somata and **C** dendrites of motor neurons expressing ALS variants compared to eGFP control. **D,E** Reduction in *Hspa8* mRNA only in somata of neurons expressing wild type (WT) TDP-43. Data presented as mean spot counts ± S.D., n = 11-61 neurons. **F,G** Reduction in HSPA8 protein in somata of motor neurons expressing ALS variants, compared to noninjected motor neurons in the same culture (CTL). **F** Representative micrographs of spinal cord-DRG culture double-labeled with anti-HSPA8 and anti-TDP-43. Arrow points to a neuron expressing TDP-43^G348C^. **G** Graphed is mean pixel intensity of immunolabeling signal ± S.D., n = 9-24 neurons. Statistical significance was evaluated through one-way ANOVA followed by Bonferroni post hoc analysis. **p<0.01, ***p<0.001, ****p<0.0001. Scale bar = 15μm.

The reduction in *Hspa8* mRNA levels in somata of motor neurons expressing ALS variants was reflected in the decreased expression of HSPA8 protein (Fig. 3F,G).

#### Impact of RGFP963 and arimoclomol on *Hspa8* mRNA levels in motor neurons expressing ALS variants

Only combined treatment with RGFP963 and arimoclomol significantly maintained *Hspa8* mRNA counts in somata of motor neurons expressing TDP-43^G348C^ – mean count of 1340 mRNA spots compared to 590 with vehicle treatment, on par with the control levels (Fig. 4A). In contrast, drug treatments failed to maintain dendritic *Hspa8* mRNA levels due to retention in somata (Fig. 4B).

**Figure 4.**
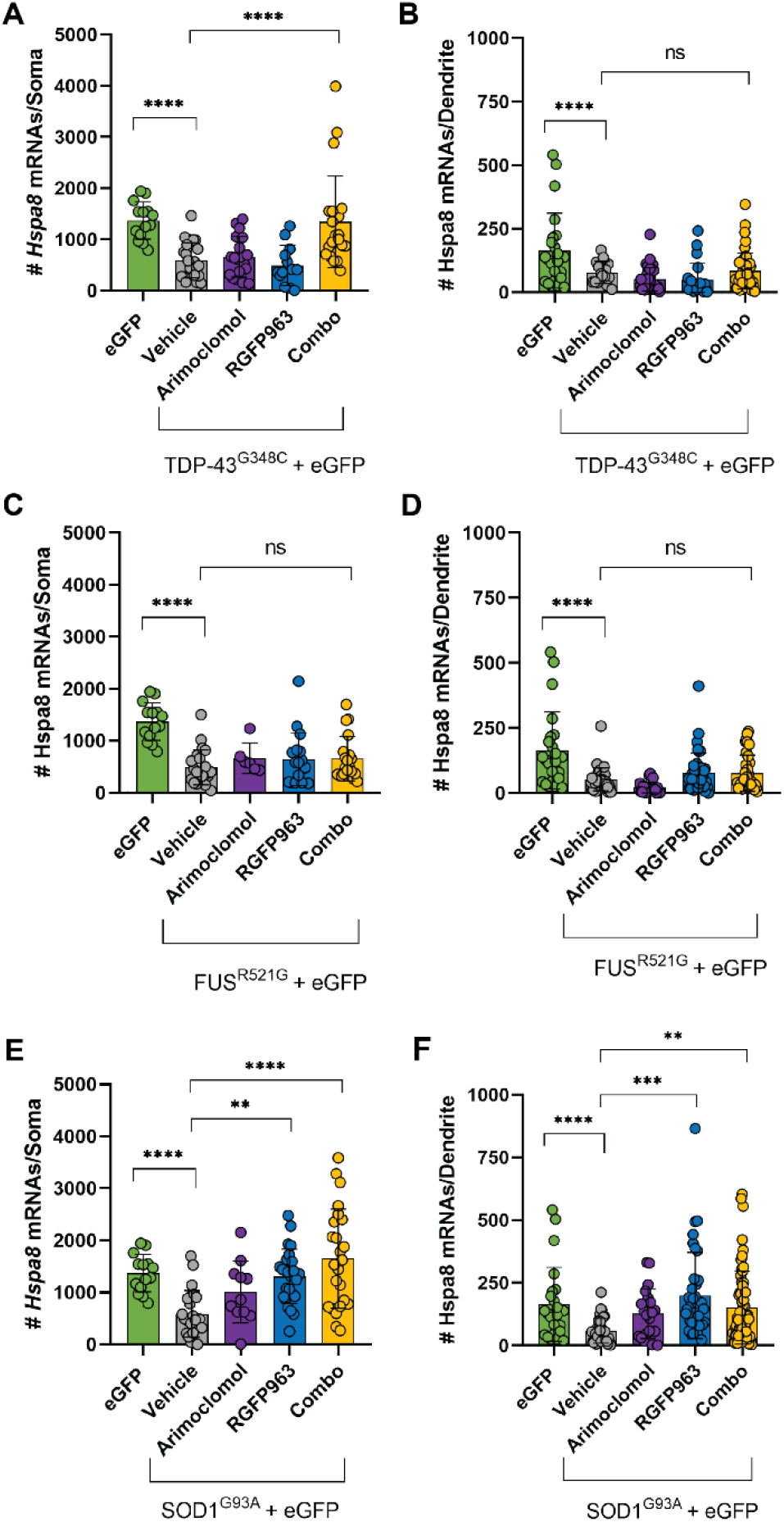
Effect of RGFP963 and arimoclomol alone and in combination on *Hspa8* mRNA levels in motor neurons expressing ALS variants. smFISH quantitation of *Hspa8* mRNA levels in both somata and dendrites of motor neurons expressing eGFP alone (control) or with the ALS-associated variants TDP-43^G348C^, FUS^R521G^ or SOD1^G93A^. Cultures were treated with vehicle (DMSO) or with 4 µM arimoclomol or 1µM RGFP963 alone or in combination. *Hspa8* smFISH and immunolabeling with antibodies to eGFP and TDP-43, FUS or SOD1 was carried out on day three. **A,B TD-43 model:** Only the combination of RGFP963 and arimoclomol preserved *Hspa8* mRNA levels in somata. Levels remained low In dendrites, regardless of the treatment. **C,D FUS^R521G^ model:** Neither individual treatments nor their combination maintained the levels of *Hspa8* mRNA in either somata or dendrites. **E,F SOD1 model:** Both RGFP963 alone and combined with arimoclomol, but not arimoclomol alone, significantly maintained *Hspa8* mRNA levels in both somata and dendrites. Data are presented as mean ± S.D., n = 14-60 neurons. Statistical significance was evaluated through one-way ANOVA followed by Bonferroni post hoc analysis. **p<0.01, ***p<0.001, ****p<0.0001.

In motor neurons expressing FUS^R521G^, neither individual treatments nor their combination maintained the levels of *Hspa8* mRNA in somata or dendrites (Fig. 4C,D). In contrast, in motor neurons expressing SOD1^G93A^, both individual treatment with RGFP963 and its combination with arimoclomol effectively maintained *Hspa8* mRNA levels in both the somata and dendrites at near control levels (Fig. 4E,F). Arimoclomol treatment was ineffective in maintaining dendritic *Hspa8* mRNA, although there was a trend to maintain levels in the somata.

#### Longitudinal analysis of *Hspa8* mRNA levels in motor neurons expressing ALS variants

To detect variations in mRNA levels within the initial days of SOD1^G93A^, TDP-43^G348C^, or FUS^R521G^ expression, we quantified *Hspa8* mRNA over a three-day period, to capture any initial but transient changes that might have resolved by day three.

*Hspa8* mRNA levels in both non-injected and eGFP-expressing motor neurons remained consistent over the course of three days (Fig. 5A,B). In somata of motor neurons expressing TDP-43^G348C^, *Hspa8* mRNA levels were elevated on day one (mean of 3000 mRNAs) and progressively declined over days two and three (mean of 600 mRNAs) (Fig. 5C). Again, treatment with the combination of RGFP963 and arimoclomol prevented the decline on day three, maintaining mean count close to control (mean of 1300 mRNAs) (Fig. 5C), but had no effect on dendritic levels (Fig. 5D).

**Figure 5.**
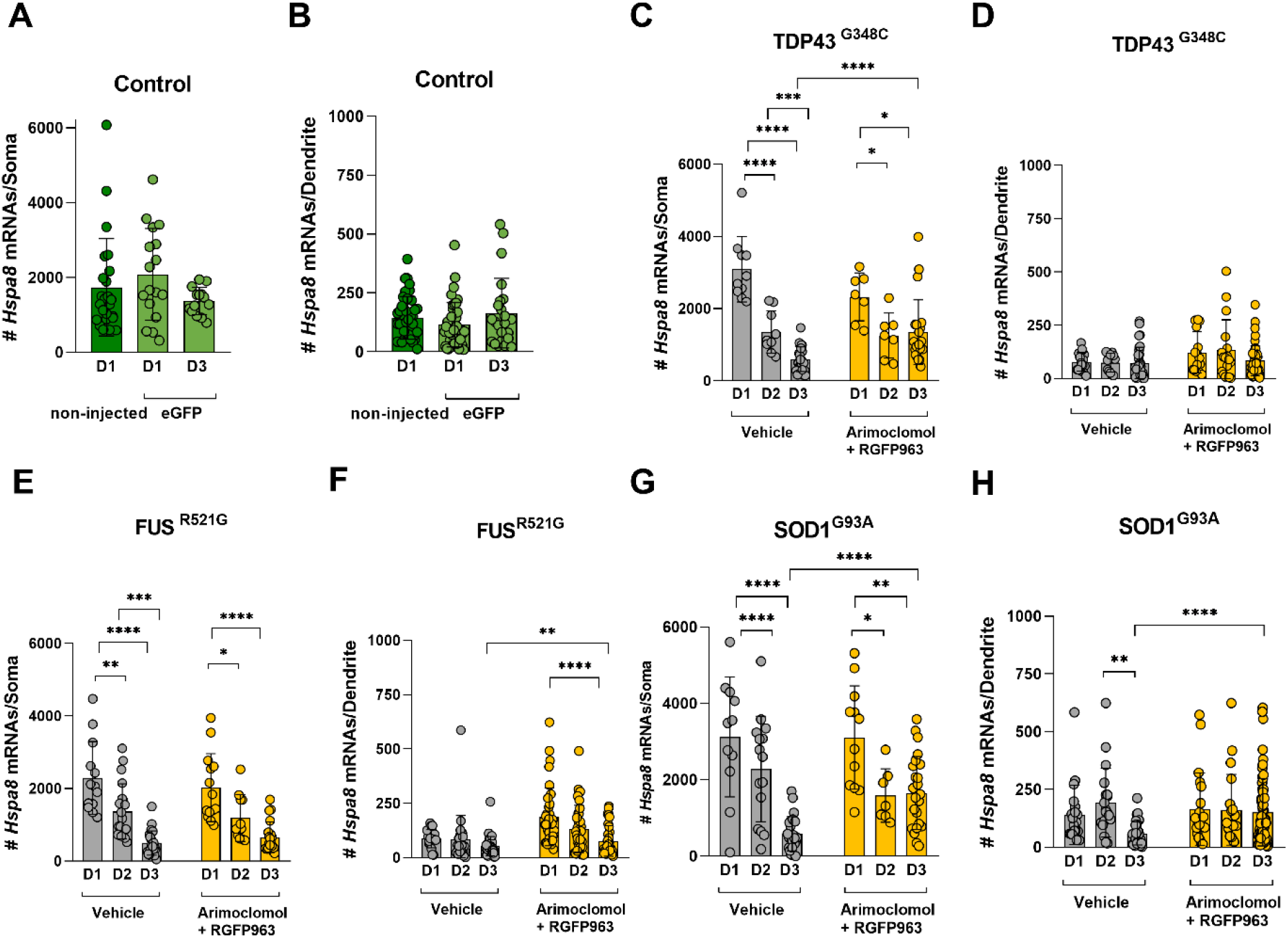
Differential effects of arimoclomol and RGFP963 on the decline in *Hspa8* mRNA levels in motor neurons expressing ALS variants. **A,B** Controls: *Hspa8* mRNA expression remained unchanged over a three-day period in **A** soma and **B** dendrites of control motor neurons (non-microinjected neurons labeled with MAP2 antibody and microinjected motor neurons expressing eGFP). **C,D** In somata of motor neurons expressing TDP-43^G348C^, *Hspa8* levels gradually declined over three days; Combined treatment with RGFP963 and arimoclomol prevented the the decline from day two to day three. **E,F** In motor neurons expressing FUS^R521G^, *Hspa8* mRNA levels declined in both somata and dendrites. Combination treatment failed to prevent this decline in somata, although decline in dendrites was less. **G,H** Spike in *HSPA8* mRNA somata of SOD1^G93A^-expressing motor neurons on day one, but substantial decline by day three, which was prevented by the drug combination. Similarly, drug combination maintained *Hspa8* mRNA in dendrites. Data are presented as mean ± S.D., n = 9-60 neurons. Statistical significance was evaluated through one-way ANOVA followed by Bonferroni post hoc analysis. **p<0.01, ***p<0.001, ****p<0.0001.

In FUS^R521G^-expressing motor neurons there was also a progressive decline in the mean number of *Hspa8* mRNA spots over the three-day period, from 2000 on day one, to 1300 on day two and to 490 on day three. Combined treatment with RGFP963 and arimoclomol had no effect (Fig. 5E). A similar gradual decline in dendritic *Hspa8* mRNA levels occurred in dendrites, although in this experiment there was some preservation on day 3 by the combination treatment (Fig. 5F).

In motor neurons expressing SOD1^G93A^, *Hspa8* mRNA levels in somata were initially increased (day one mean of 3100 mRNAs in comparison to 1800 in eGFP control, P= 0.0101) (Fig. 5G), but subsequently declined to 2200 mRNAs on day two and only 580 mRNAs by day three. Combined treatment with RGFP963 and arimoclomol prevented the decline between day two and day three. Similarly in dendrites, expression of SOD1^G93A^ reduced *Hspa8* mRNA levels over the three days, which was prevented by combination treatment (Fig. 5H).

In summary, ALS variants were poor inducers of *Hspa1a* expression; RGFP963 slightly improved this response in motor neurons expressing TDP-43 and SOD1 variants, but arimoclomol was significantly effective only in the SOD1 model. *Hspa8* mRNA and HSPA8 protein levels were robust in both somata and dendrites of motor neurons but declined over three days with expression of TDP-43^G348C^, FUS^R521G^ or SOD1^G93A^. Arimoclomol had minimum impact on these measures, whereas RGFP963 was particularly effective in maintaining *Hspa8* expression in the SOD1 model. Whether this preservation involved specific action on *Hspa8* gene transcription by the drugs or occurred secondary to other drug actions that preserved motor neuron functions more generally would be difficult to determine. Nevertheless, the previously reported neuroprotection in the FUS model by HDAC inhibitors and arimoclomol, in the absence of any measured upregulation of HSPA1A or HSPA8, encouraged us to measure biomarkers of toxicity in the TDP-43 model.

### Impact of RGFP963, arimoclomol alone and in combination on disease-relevant biomarkers in motor neurons expressing TDP-43^G348C^

#### Histone acetylation

There is evidence of decreased histone acetylation in ALS tissue and experimental models [13,31–35]. We previously reported that H3/H4 histone acetylation, associated with active chromatin, was reduced in cultured motor neurons expressing a fALS-linked FUS variant and was prevented by treatment with the pan HDAC inhibitor, SAHA [35]. Thus, examining the effect of HDAC inhibitors serves as a biomarker of target engagement as well as addressing a relevant biomarker of disease.

To assess histone acetylation in motor neurons expressing TDP-43^G348C^, we used an antibody specifically recognizing H3 acetylated at residues 9 and 14 (H3K9/K14Ac) and rated antibody labeling of the nucleus as strong (H3K9Ac+/+), weak (H3K9Ac+/-) or absent (H3K9Ac-/-) (Fig. 6A). In line with our previous findings [36], we observed decreased H3K9/K14 acetylation in motor neurons expressing TDP-43^G348C^. Sixty percent of control motor neurons exhibited strong nuclear H3K9/K14Ac+/+ compared to 28% of TDP-43^G348C^-expressing motor neurons (Fig. 6B). This percentage was significantly increased to control levels by treatment with SAHA (Fig. 6C) or the Class I inhibitor, RGFP963 (Fig. 6D), but arimoclomol was ineffective (Fig. 5C,D), consistent with its lack of class I HDAC inhibitory activity [7]. Expression of wild-type TDP-43 did not significantly alter H3K9/K14Ac scoring (Fig. 6E), nor was H3 acetylation influenced by SAHA (Fig. 6E).

**Figure 6.**
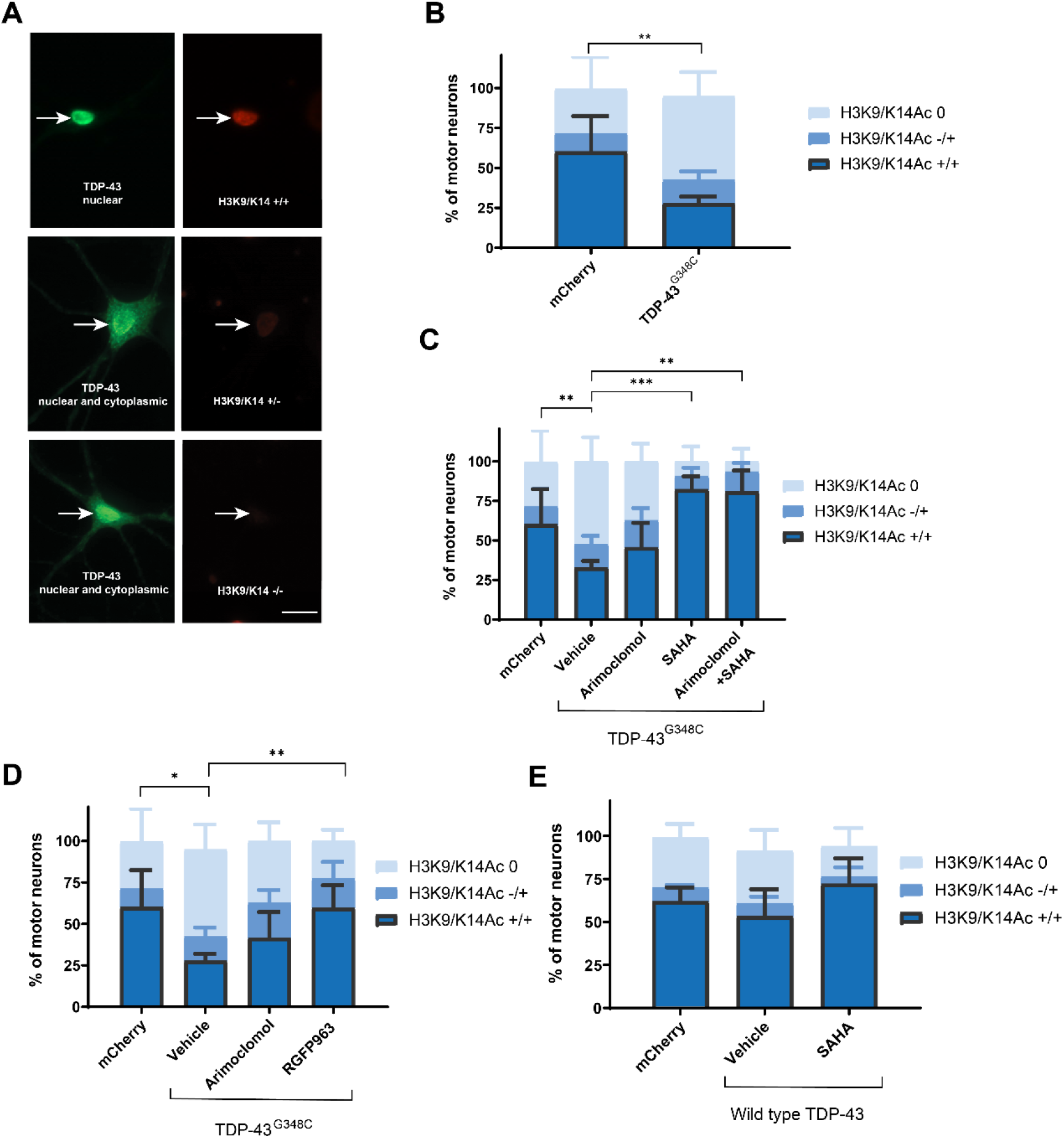
HDAC inhibition limited the reduction of histone acetylation in TDP-43^G348C^-expressing motor neurons. After microinjection of plasmid encoding TDP-43^G348C^ or mCherry as control, cultures were treated with vehicle (DMSO), 4 µM arimoclomol, 4µM SAHA, 1µM RGFP963, alone and in combination for three days. Injected neurons were identified by double-labeling with rabbit anti-flag M2 to visualize TDP-43^G348C^ or by mCherry epifluorescence. **A** Acetylation of histone 3 at lysine residues 9 and 14 (H3K9/K14Ac), evaluated by immunocytochemistry using an acetylation-specific antibody was categorized as: strong (H3K9/K14Ac+/+), weak (H3K9/K14Ac+/-) or absent (H3K9/K14Ac-/-). **B** Reduction in H3K9K14ac in motor neurons expressing TDP-43^G348C^ was prevented by the HDAC inhibitors **C** SAHA or **D** RGFP963 demonstrating target engagement, whereas arimoclomol had no impact. **E** No alteration of histone acetylation by wild-type TDP-43. Data are presented as mean ± S.D., n = 6-13 cultures. Statistical significance was evaluated by one-way ANOVA followed by Bonferroni post hoc analysis. *p<0.05, **p<0.01, ***p<0.001. Scale bar = 20µm.

#### Nuclear-cytoplasmic localization of TDP-43

Abnormal cytoplasmic accumulation of RNA-binding proteins is a hallmark of ALS, potentially contributing to nuclear dysfunction and cytoplasmic toxicity [37]. This feature is reproduced in cultured motor neurons expressing ALS-linked TDP-43 variants [25] and FUS variants [15]. We previously reported that SAHA and RGFP109 maintained FUS within the nucleus of cultured motor neurons expressing FUS^R521H^, increasing the percentage of neurons exhibiting both nuclear and cytoplasmic FUS and decreasing the percentage in which FUS was exclusively cytoplasmic [7]. Thus, we determined if these treatments had comparable effects on distribution of TDP-43 in motor neurons expressing TDP-43^G348C^.

Three days following microinjection of expression plasmid, cultures were immunolabeled with antibody to TDP-43 and immunolabeling was categorized as: cytoplasmic only, both nuclear and cytoplasmic, and nuclear only (Fig. 7A). Treatment with SAHA (Fig. 7B), RGFP963 (Fig. 7C), or RGFP109 (Fig.7D) significantly preserved nuclear TDP-43, with a significantly lower percentage of neurons expressing the variant scored as having only cytoplasmic TDP-43. A similar preservation of nuclear TDP-43 was achieved by treatment with arimoclomol. As reported in the FUS ALS model [7], the HDAC6 inhibitor Tubastatin A had no significant effect on TDP-43 localization (Fig. 7E).

**Figure 7.**
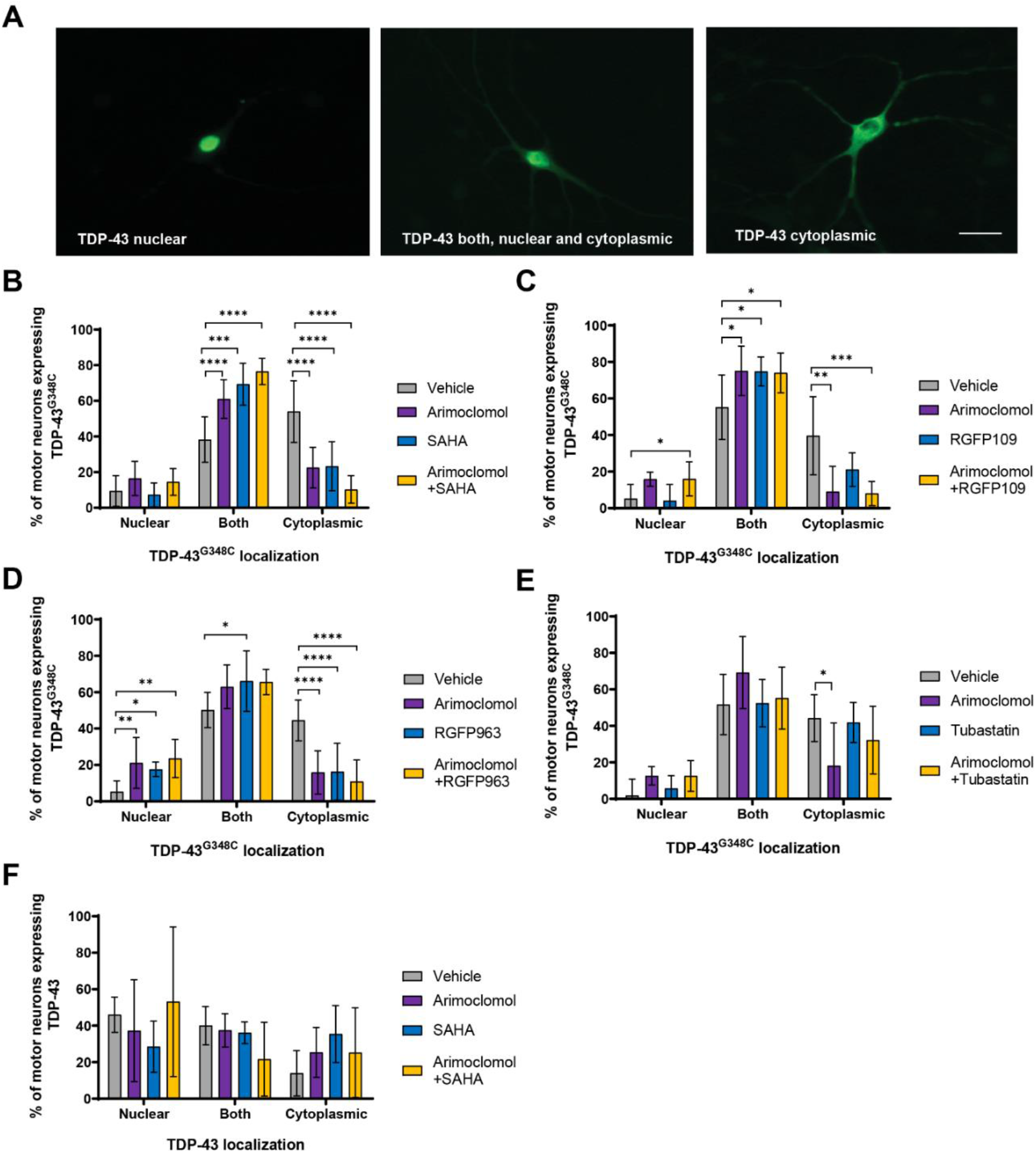
HDAC inhibitors and arimoclomol preserve nuclear localization of TDP-43. in motor neurons expressing. Cultures containing motor neurons expressing TDP-43^G348C^ or wild-type TDP-43 were treated with vehicle (DMSO), 4 µM arimoclomol, or HDAC inhibitor (4µM SAHA, 1µM RGFP963, 4µM RGFP109 or 1µM Tubastatin A) alone or in combination with arimoclomol. **A** After three days, cultures were immunolabeled with rabbit anti-TDP-43 and TDP-43 localization was categorized as: cytoplasmic only, both nuclear and cytoplasmic, and nuclear only. **B** SAHA, **C** RGFP109, or **D** RGFP963 significantly preserved nuclear TDP-43 in motor neurons expressing TDP-43^G348C^. Arimoclomol treatment also significantly maintained nuclear TDP-43. **E** No effect of the HDAC6 inhibitor Tubastatin A. **F** Neither overexpression of wild-type TDP-43 nor treatment of these neurons with drugs affected TDP-43 distribution. Data are presented as mean ± S.D. n = 6-9 cultures. Statistical significance was evaluated by one-way ANOVA followed by Dunnett’s multiple comparison test. *p<0.05, **p<0.01, ***p<0.001, ****p<0.0001. Scale bar = 20µm.

Some motor neurons overexpressing wild-type TDP-43 lacked nuclear TDP-43 (Fig. 7F), consistent with previous data and the normal cycling of this RNA-binding protein between the nucleus and cytoplasm in neurons [25]; however, neither arimoclomol nor SAHA treatment significantly affected wild type TDP-43 localization.

#### Expression of Brg1, the nBAF chromatin remodeling complex ATPase

Components of the nBAF chromatin-remodeling complex, including the ATPase Brg1, are depleted from the nucleus of cultured motor neurons expressing ALS-linked FUS or TDP-43 variants, correlated with reduction in nuclear FUS or TDP-43 and with dendritic attrition [13]. nBAF complex depletion also occurred in spinal motor neurons in autopic tissue from both sALS and fALS ALS cases [13]. We now report similar findings in cultured motor neurons expressing TDP-43^G348C^. Nuclear Brg1 was evaluated by immunolabeling in the absence and presence of treatment with HDAC inhibitor (SAHA or RGFP963), arimoclomol alone, or drug combinations. Immunolabeling with antibody to Brg1 was categorized based on labeling intensities: strong (+/+), weak (+/-), or none (-/-) as shown in Fig. 8A. The mean percentage of TDP-43^G348C^-expressing motor neurons scored as (+/+) Brg1 was reduced to 35% compared to 62% in the control group (expressing mCherry) (Fig. 8B). Both SAHA (Fig. 8C) and RGFP963 (Fig. 8D) maintained nuclear Brg1 at levels comparable to control motor neurons, 65% and 61% of motor neurons, respectively scoring as (+/+) Brg1. Arimoclomol also preserved nuclear Brg1 within TDP-43^G348C^-expressing motor neurons (mean (+/+) score of 57%). Overexpression of wild-type TDP-43 did not significantly affect nuclear Brg1 scoring (Fig. 8E).

**Figure 8.**
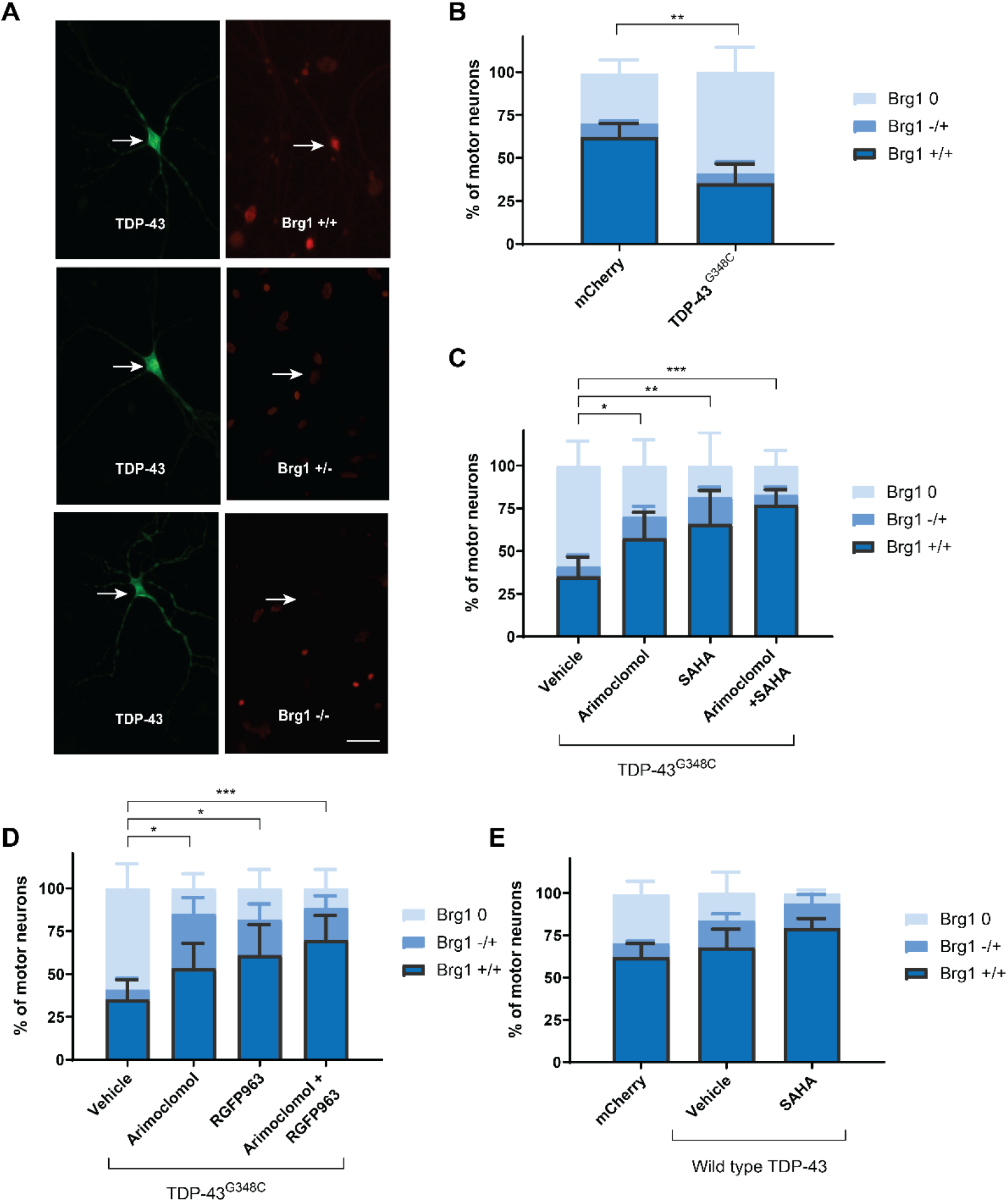
HDAC inhibitors and arimoclomol preserve Brg1, the ATPase component of nBAF chromatin remodeling complexes, in TDP-43^G348C^-expressing motor neurons. Three days after microinjection, cultures were immunolabeled with rabbit anti-Brg1 and mouse anti-flag M2 to visualize TDP-43^G348C^. **A** Intensity of Brg1 immunolabeling in the nucleus was categorized as: strong (+/+), (+/-), or none (-/-). **B** Compared to controls (mCherry), expression of TDP-43^G348C^ significantly reduced the percentage of motor neurons displaying strong Brg1 labeling. This percentage was maintained similar to controls by **C** SAHA and **D** RGFP963 treatments. Arimoclomol also maintained nuclear Brg1 expression, although slightly less efficiently than HDAC inhibitors. When SAHA or RGFP963 was combined with arimoclomol, a modest enhancement occurred. **E** Brg1 distribution was not significantly affected by over-expression of wild-type TDP-43. Data are presented as mean ± S.D. n = 4-15 cultures. Statistical significance was evaluated through one-way ANOVA followed by Bonferroni post hoc analysis. *p<0.05, **p<0.01, ***p<0.001. Scale bar = 20µm.

### Effect of fALS variants, HDAC inhibition and arimoclomol on axonal transport of mitochondria

Disruption of organelle transport, including mitochondria, is a common consequence of expression of ALS-linked variants (reviewed in [38]). For vital imaging of transport in axonal segments, mCherry was targeted to mitochondria by incorporating the ornithinecarbamoyltransferase signal sequence (Fig. 9A). Fig. 9A includes sample kymograph recordings, showing color-coded mitochondria. The mean start-to-end velocity was significantly reduced by expression of TDP-43^G348C^, FUS^R521G^ or SOD1^G93A^ (Fig. 9B). Neither RGFP963 nor arimoclomol preserved transport in neurons expressing TDP-43^G348C^ (Fig. 9C) or FUS^R521G^ (Fig. 9D). In the SOD1 model, transport was maintained at control levels by treatment with RGFP963 in combination with arimoclomol, but not by these drugs individually, another example of the drug combination outperforming individual drugs (Fig. 9E).

**Figure 9.**
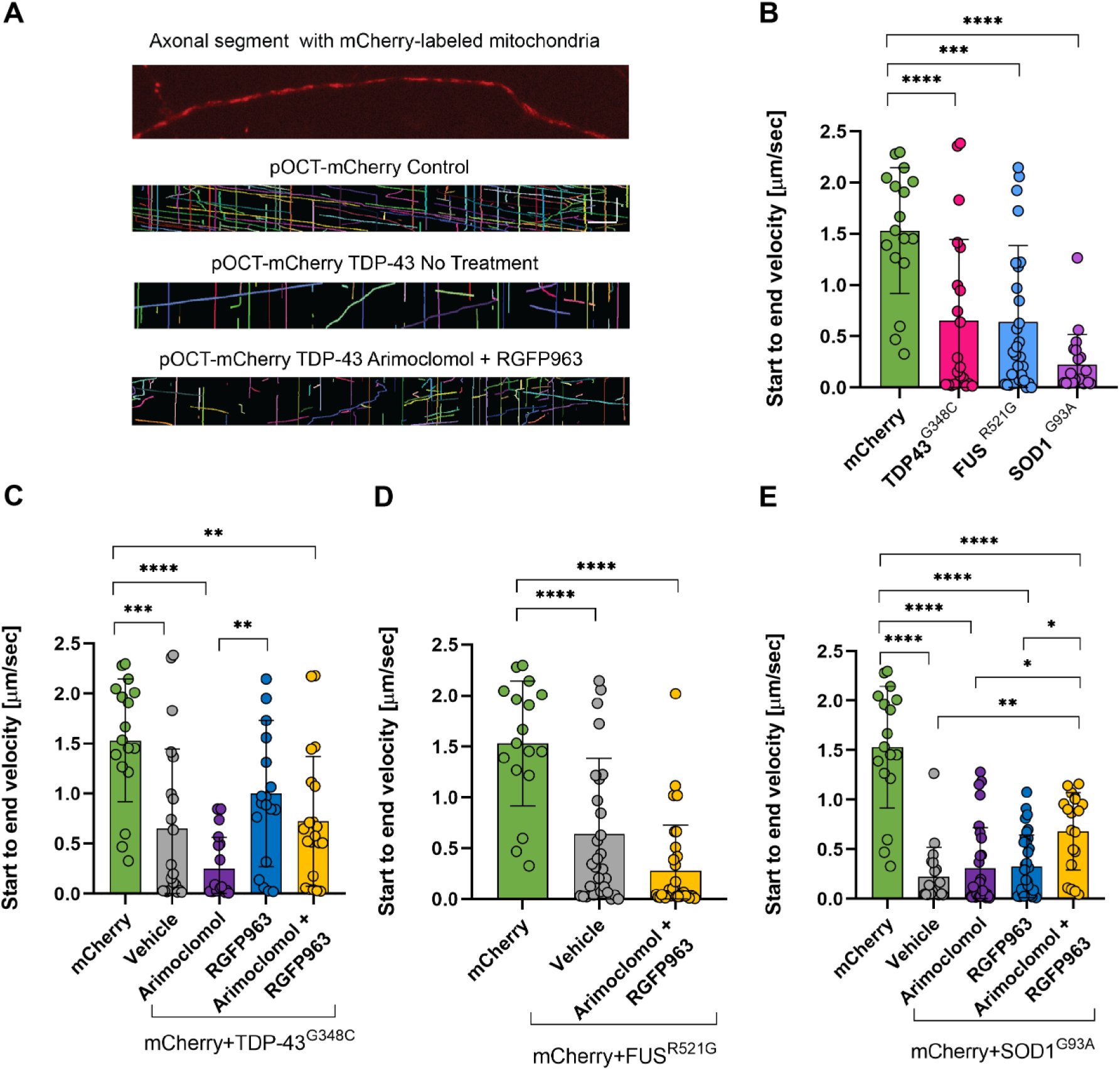
Effect of ALS variants and drug treatments on axonal transport of mitochondria. **A** Image of axonal segment of motor neuron expressing pOCT-mCherry localized to mitochondria. Kymographs generated from mitochondrial movements in control or TDP-43^G348C^-expressing neurons and in cultures treated with vehicle or the combination of arimoclomol and RGFP963. Individual mitochondria were color-coded using KymoResliceWide. Scale bars = 20µm horizontal and 3.5 min vertical. **B** TDP-43^G348C^, FUS^R521G^ or SOD1^G93A^ impaired start-to-end velocity of mitochondrial transport in axonal segments of motor neurons compared to mCherry alone. **C** Neither RGFP963 nor arimoclomol preserved mitochondrial transport in neurons expressing TD-P43^G348C^ or **D** FUS^R521G^. Only the combination of RGFP963 and arimoclomol reduced the decline in mitochondrial transport by SOD1^G93A^. Data are presented as mean ± S.D. n = 17-33 axons per group. Statistical significance was evaluated through one-way ANOVA followed by Bonferroni post hoc analysis. *p<0.05, **p<0.01, ***p<0.001.

## Discussion

ALS is a heterogeneous disease, characterized by multiple interacting pathophysiological mechanisms that can vary with genetic factors, environmental influences and stage of disease. Although protein quality control mechanisms have been recognized as a potential cross-cutting therapeutic strategy, they have proven difficult to effectively manipulate for clinical benefit. Using primary culture models of three forms of ALS caused by mutations in *TARDPB*, *FUS* and *SOD1*, we have highlighted the differences in the effect of these variants on expression of HSPs, specifically the stress-inducible isoform HSPA1A and the constitutively expressed HSC HSPA8, as well as the effect of drugs that potentially alter their expression. This paper focused on the *TARDBP* model to complement a previous study using *SOD1* and *FUS* models [7], as well as on additional experiments comparing all three models (summarized in Table 1 in Supplementary data).

Poor induction of HSPA1A with proteotoxic stress has remained a consistent theme, with FUS variants being particularly ineffective [2,7]. Several factors have been proposed to underly the lack of HSPA1A expression in neurons. Levels of the transcription factor HSF1 are low in neurons and generally diminish with aging [9], but overexpression of wild type HSF1 in cultured motor neurons was not sufficient [39]. This study demonstrated the lack of *Hspa1a* mRNA expression as well. TDP-43 and FUS variants affect multiple levels of gene and protein expression including RNA splicing, transport and translation that could contribute [37,40–43], but we observed transcription sites appearing only transiently, if at all, in motor neurons expressing ALS variants, pointing to the major impairment in HSPA1A induction being at the transcriptional level.

Translation of *Hspa1a* mRNA requires the translation initiation factor eEF1A1, but the neuron-specific variant eEF1A2 lacks this ability [44]. Expression switches from eEF1A1 to eEF1A2 during neuronal development, although some eEF1A1 has been localized to postsynaptic densities [45,46] and axons [47], which could support local translation. In Drosophila brain, Mettl3-dependent increase in m^6^ A (methyladenosine) modification of polyA+ RNA with heat stress attenuated the heat shock response at the translational level [48]. *Hspa1a* mRNA was very low in motor neurons expressing ALS variants arguing against the major block being at the translational stage. Nevertheless, *Hspa1a* transcription can be increased – two conditions that do upregulate HSPA1A in cultured motor neurons are treatment with agents that disrupt HSP90 complexes that sequester HSF1 [2,8] and proteasome inhibition, which leads to increased levels of the heat shock transcription factor HSF2 [2].

We postulated that HDAC inhibitors by promoting histone acetylation and openness of chromatin would facilitate access of HSF1 to HSE of heat shock genes and improve the efficacy of HSP co-inducers, such as arimoclomol, when used in combination. HDAC inhibition did preserve histone 3 acetylation in motor neurons expressing FUS [35] or TDP-43 variants (this study). While class I HDAC inhibitors were as effective as arimoclomol in inducing HSPA1A expression under various stress conditions [7], in only some circumstances has the combination of HDAC inhibitor with arimoclomol been more effective than either drug alone; *e.g*., more motor neurons expressing SOD1^G93A^ were immunopositive for HSPA1A when treated with SAHA and arimoclomol in combination compared to either drug alone [7]. However, in motor neurons expressing TDP-43^G348^, arimoclomol was not effective, nor did combining with HDAC inhibitors improve efficacy. Neurons expressing FUS^R521G^ failed to respond to any treatment by expressing HSPA1A at the protein or mRNA level.

There is increasing appreciation for epigenetic mechanisms contributing to ALS and other neurodegenerative disorders including DNA methylation, chromatin remodeling factors and post-translational modification of histones [3,13,49–51]. Recruitment of chromatin remodeling complexes by HSF1 is an important aspect of transcriptional regulation [52]; thus, the loss of nBAF chromatin remodeling complexes in ALS might be disruptive. Both class I HDAC inhibition and arimoclomol preserved nuclear TDP-43, FUS and Brg1 (the ATPase component of nBAF chromatin remodeling complexes) in motor neurons, but without promoting *Hspa1a* expression.

Several families of HSPs cooperate for protein quality control and might obviate a role for HSPA1A under real world conditions in mammalian systems. Expression of the HSC, HSPA8, was suppressed in both somata and dendrites in all three ALS models tested, both at the mRNA and protein level, which would likely have significant impact on proteostasis. HSPA8 interacts with TDP-43 [53,54]. In Drosophila motor neurons, TDP-43^G298S^ interacted with Hsc70-4 (HSPA8) protein and mRNA to a greater extent than wild type TDP-43 and suppressed its translation, with reduction in Hsc70-4 levels at neuromuscular junction synapses [55]. HSPA8 also was reduced in growth cones of primary cultured motor neurons overexpressing TDP-43^Q331K^ or TDP-43^M337V^ and at neuromuscular junctions of TDP-43^A315T^ transgenic mice, with impairment of synaptic vesicle endocytosis. In our study, the suppression of HSPA8 was observed in soma, perhaps due to higher level of overexpression of TDP-43^G348C^ compared to models used by Coyne et al., but the message is qualitatively similar.

RGFP963 preserved *Hspa8* mRNA in somata and dendrites of motor neurons expressing SOD1^G93A^, but only in somata of neurons expressing TDP-43^G348C^ when combined with arimoclomol. No drug treatment was effective in the FUS ALS model, following the same pattern as with HSPA1A. The distinction between somatic and dendritic mRNA levels is significant given that *Hspa8* mRNA is the chaperone mRNA most transported to dendrites for translation during proteotoxic stress and this transport is impaired with knockdown of FUS or expression of a dominant mutation in HNRNPA2B1, another RNA-binding protein [30]. Thus, RNA binding proteins play an important role in dendritic proteostasis. To maintain neuronal connectivity, an effective therapy would need to not only regulate gene expression but target proteins to appropriate cellular compartments.

As in Kuta et al. (2020), efficacy of HDAC inhibition, arimoclomol and the combination depended on the stress and the biomarker being measured. Indeed, arimoclomol presents an enigma. It is inherently inefficient as an HSP co-inducing agent, yet it protects against some measures of toxicity in ALS models, including the FUS model, which was the least responsive in terms of HSPA1A induction and preservation of *Hspa8* mRNA expression. Indeed, arimoclomol failed to maintain *Hspa8* mRNA expression in neurons expressing ALS variants, despite preserving nuclear TDP-43 and FUS and nBAF chromatin remodeling complexes (this study and Kuta et al. (2020)). On the other hand, HDAC inhibition, but not arimoclomol, inhibited cytoplasmic aggregation FUS in cultured motor neurons expressing FUS^R521G^, contrary to what might be expected of a HSP coinducer (Supplementary Fig. 2).

Neither arimoclomol nor RGFP963 significantly preserved mitochondrial transport in motor neurons expressing TDP-43^G348C^ or FUS^R521G^. Tubulin acetylation is an important regulator of axonal transport, but since this posttranslational modification is regulated by the Class IIB HDAC, HDAC6, Class I inhibitors would not be expected to have a direct effect on transport mechanisms [56]. In the SOD1 model, the combination of arimoclomol and RGFP963 did improve transport, although the mechanism(s) aren’t apparent. Mitochondria are particularly compromised by expression of SOD1 variants [57], including accumulating Ca^++^ and vacuolating, abnormalities that are expressed in our primary motor neuron culture model in addition to impaired transport [24].

In conclusion, the multitude of interacting pathways altered by expression of these fALS variants and in ALS in general, and incomplete understanding of drug properties, defies identifying precise mechanisms of neuroprotection. Collectively, the data speak to the complexity of drug mechanisms and the importance of testing drug combinations against multiple biomarkers of ALS pathogenesis to identify effective therapies depending on the type and stage of disease.

## Acknowledgements

This study was supported by an ALS Canada-Brain Canada Hudson Translational Team Grant (HDD, JN, CFS and RR) and ALS Discovery grant (MV and HD) provided by Brain Canada through the Canada Brain Research Fund with the financial support of Health Canada and the ALS Society of Canada. MFC was supported by Fonds de Recherche du Québec Nature et Technologies and Mitacs Globalink Graduate Fellowship, Consejo Nacional de Ciencia y Tecnología. RGFP963 was a kind gift of BioMarin Pharmaceutical Inc., San Raphael, CA USA. These funding sources had no role in the conduct of the research and/or preparation of the article.

The authors have no competing interests relevant to this project.

## REFERENCES

[1] P. Manzerra, I. R. Brown, Expression of heat shock genes (hsp70) in the rabbit spinal cord: Localization of constitutive and hyperthermia-inducible mRNA species, J. Neurosci. Res. 31 (1992) 606–615. doi: 10.1002/jnr.490310404.

[2] Z. Batulan, et al., High threshold for induction of the stress response in motor neurons is associated with failure to activate HSF1, J. Neurosci. 23 (2003) 5789–5798. https://www.ncbi.nlm.nih.gov/pmc/articles/PMC6741252/.

[3] J. Labbadia, et al., Altered chromatin architecture underlies progressive impairment of the heat shock response in mouse models of Huntington disease, J. Clin. Invest. 121 (2011) 3306–3319. 10.1172/JCI57413.

[4] J. Hargitai, et al., Bimoclomol, a heat shock protein co-inducer, acts by the prolonged activation of heat shock factor-1, Biochem. Biophys. Res. Commun. 307 (2003) 689–695. doi: 10.1016/s0006-291x(03)01254-3.

[5] D. Kieran, et al., Treatment with arimoclomol, a coinducer of heat shock proteins, delays disease progression in ALS mice, Nat. Med. 10 (2004) 402–405. 10.1038/nm1021.

[6] M. Benatar, et al., Randomized, double-blind, placebo-controlled trial of arimoclomol in rapidly progressive SOD1 ALS, Neurology. 90 (2018) doi: e565–e574. 10.1212/WNL.0000000000004960.

[7] R. Kuta, et al., Depending on the stress, histone deacetylase inhibitors act as heat shock protein co-inducers in motor neurons and potentiate arimoclomol, exerting neuroprotection through multiple mechanisms in ALS models, Cell Stress Chaperones. 25 (2020) 173–191. doi: 10.1007/s12192-019-01064-1.

[8] Z. Batulan, et al., Induction of multiple heat shock proteins and neuroprotection in a primary culture model of familial amyotrophic lateral sclerosis, Neurobiol. Dis. 24 (2006) 213–225. 10.1016/j.nbd.2006.06.017.

[9] S. W. Kmiecik, M. P. Mayer, Molecular mechanisms of heat shock factor 1 regulation, Trends Biochem Sci. 47 (2022) 218–234. doi: 10.1016/j.tibs.2021.10.004.

[10] E. K. Sullivan, et al., Transcriptional activation domains of human heat shock factor 1 recruit human SWI/SNF, Mol. Cell Biol. 21 (2001) 5826–5837. doi: 10.1128/mcb.21.17.5826-5837.2001.

[11] J. I. Wu, et al., Regulation of dendritic development by neuron-specific chromatin remodeling complexes, Neuron. 56 (2007) 94–108. 10.1016/j.neuron.2007.08.021.

[12] I. Nakano, A. Hirano, Atrophic cell processes of large motor neurons in the anterior horn in amyotrophic lateral sclerosis: observation with silver impregnation method., J. Neuropathol. Exp. Neurol. 46 (1987) 40–49. doi: 10.1097/00005072-198701000-00004.

[13] M. Tibshirani, et al., Dysregulation of chromatin remodelling complexes in amyotrophic lateral sclerosis, Hum Mol Genet. 26 (2017) 4142–4152. Doi: 10.1093/hmg/ddx301.

[14] C. Vance, et al., Mutations in FUS, an RNA processing protein, cause familial amyotrophic lateral sclerosis type 6, Science. 323 (2009) 1208–1211. 10.1126/science.1165942.

[15] M. L. Tradewell, et al., Arginine methylation by PRMT1 regulates nuclear-cytoplasmic localization and toxicity of FUS/TLS harbouring ALS-linked mutations, Hum. Mol. Genet. 21 (2012) 136–149. 10.1093/hmg/ddx301.

[16] V. M. Van Deerlin, et al., TARDBP mutations in amyotrophic lateral sclerosis with TDP-43 neuropathology: a genetic and histopathological analysis, Lancet Neurol. 7 (2008) 409–416.

[17] E. L. Scotter, et al., TDP-43 Proteinopathy and ALS: Insights into Disease Mechanisms and Therapeutic Targets, Neurotherapeutics. 12 (2015) 352–363. doi: 10.1007/s13311-015-0338-x.

[18] I. R. Mackenzie, et al., Pathological TDP-43 distinguishes sporadic amyotrophic lateral sclerosis from amyotrophic lateral sclerosis with SOD1 mutations, Ann. Neurol. 61 (2007) 427–434. doi: 10.1002/ana.21147.

[19] M. Neumann, et al., Ubiquitinated TDP-43 in frontotemporal lobar degeneration and amyotrophic lateral sclerosis, Science. 314 (2006) 130–133. doi: 10.1126/science.1134108.

[20] M. Neumann, et al., Ubiquitinated TDP-43 in frontotemporal lobar degeneration and amyotrophic lateral sclerosis, Science. 314 (2006) 130–133. doi: 10.1126/science.1134108.

[21] S. Chen, I. R. Brown, Neuronal expression of constitutive heat shock proteins: implications for neurodegenerative diseases, Cell Stress. Chaperones. 12 (2007) 51–58. doi: 10.1379/csc-236r.1.

[22] J. Roy, et al., Glutamate potentiates the toxicity of mutant Cu/Zn-superoxide dismutase in motor neurons by postsynaptic calcium-dependent mechanisms, J. Neurosci. 18 (1998) 9673–9684. doi: 10.1523/JNEUROSCI.18-23-09673.1998.

[23] H. D. Durham, et al., Aggregation of mutant Cu/Zn superoxide dismutase proteins in a culture model of ALS, J. Neuropathol. Exp. Neurol. 56 (1997) 523–530. 10.1097/00005072-199705000-00008.

[24] M. L. Tradewell, et al., Calcium dysregulation, mitochondrial pathology and protein aggregation in a culture model of amyotrophic lateral sclerosis: Mechanistic relationship and differential sensitivity to intervention, Neurobiol. Dis. 42 (2011) 265–275. doi: 10.1016/j.nbd.2011.01.016.

[25] E. Kabashi, et al., Gain and loss of function of ALS-related mutations of TARDBP (TDP-43) cause motor deficits in vivo, Hum. Mol. Genet. 19 (2010) 671–683. doi: 10.1093/hmg/ddp534.

[26] S. Jacob-Tomas, et al., Using Single-Molecule Fluorescence Microscopy to Uncover Neuronal Vulnerability to Protein Damage, Methods Mol Biol. 2515 (2022) 237–254. doi: 10.1007/978-1-0716-2409-8_15.

[27] C. Eliscovich, et al., Imaging mRNA and protein interactions within neurons, Proc Natl Acad Sci U S A. 114 (2017) E1875–e1884. doi: 10.1073/pnas.1621440114.

[28] Z. Wefers, et al., Analysis of the Expression and Subcellular Distribution of eEF1A1 and eEF1A2 mRNAs during Neurodevelopment, Cells. 11 (2022) doi: 10.3390/cells11121877.

[29] M. A. H. Jakobs, et al., KymoButler, a deep learning software for automated kymograph analysis, Elife. 8 (2019) 10.7554/eLife.42288.

[30] C. Alecki, et al., Localized synthesis of molecular chaperones sustains neuronal proteostasis, bioRxiv. (2023) doi: 10.1101/2023.10.03.560761.

[31] C. Valle, et al., Tissue-specific deregulation of selected HDACs characterizes ALS progression in mouse models: pharmacological characterization of SIRT1 and SIRT2 pathways, Cell Death Dis. 5 (2014) e1296. doi: 10.1038/cddis.2014.247.

[32] C. Rouaux, et al., Critical loss of CBP/p300 histone acetylase activity by caspase-6 during neurodegeneration, EMBO J. 22 (2003) 6537–6549. doi: 10.1093/emboj/cdg615.

[33] C. Rouaux, et al., Sodium valproate exerts neuroprotective effects in vivo through CREB-binding protein-dependent mechanisms but does not improve survival in an amyotrophic lateral sclerosis mouse model, J. Neurosci. 27 (2007) 5535–5545. doi: 10.1523/JNEUROSCI.1139-07.2007.

[34] E. Rossaert, et al., Restoration of histone acetylation ameliorates disease and metabolic abnormalities in a FUS mouse model, Acta Neuropathol Commun. 7 (2019) 107. doi: 10.1186/s40478-019-0750-2.

[35] M. Tibshirani, et al., Cytoplasmic sequestration of FUS/TLS associated with ALS alters histone marks through loss of nuclear protein arginine methyltransferase 1, Hum. Mol. Genet. 24 (2015) 773–786. 10.1093/hmg/ddu494.

[36] M. Tibshirani, Convergent epigenetic consequences of mislocalization of ALS-linked RNA-binding proteins, Doctor of Philosophy (2016) Integrated Program in Neuroscience, McGill University, Montreal.

[37] L. Vanden Broeck, et al., TDP-43-mediated neurodegeneration: towards a loss-of-function hypothesis?, Trends Mol Med. 20 (2014) 66–71. 10.1016/j.molmed.2013.11.003.

[38] E. F. Smith, et al., The role of mitochondria in amyotrophic lateral sclerosis, Neurosci Lett. 710 (2019) doi: 132933. 10.1016/j.neulet.2017.06.052.

[39] D. M. Taylor, et al., Characterizing the role of Hsp90 in production of heat shock proteins in motor neurons reveals a suppressive effect of wild-type Hsf1, Cell Stress. Chaperones. 12 (2007) 151–162. doi: 10.1379/csc-254r.1.

[40] D. Dormann, C. Haass, TDP-43 and FUS: a nuclear affair, Trends Neurosci. 34 (2011) 339–348. doi: 10.1016/j.tins.2011.05.002.

[41] H. Deng, et al., The role of FUS gene variants in neurodegenerative diseases, Nat. Rev. Neurol. 10 (2014) 337–348. doi: 10.1038/nrneurol.2014.78.

[42] C. Lagier-Tourenne, et al., TDP-43 and FUS/TLS: emerging roles in RNA processing and neurodegeneration, Hum. Mol. Genet. 19 (2010) R46–R64. doi: 10.1093/hmg/ddq137.

[43] R. T. Bjork, et al., Dysregulation of Translation in TDP-43 Proteinopathies: Deficits in the RNA Supply Chain and Local Protein Production, Front Neurosci. 16 (2022) 840357. https://10.3389/fnins.2022.840357.

[44] M. Vera, et al., The translation elongation factor eEF1A1 couples transcription to translation during heat shock response, Elife. 3 (2014) e03164. 10.7554/eLife.03164.

[45] S. J. Cho, et al., Presence of translation elongation factor-1A (eEF1A) in the excitatory postsynaptic density of rat cerebral cortex, Neurosci Lett. 366 (2004) 29–33. doi: 10.1016/j.neulet.2004.05.036.

[46] S. J. Cho, et al., Translation elongation factor-1A1 (eEF1A1) localizes to the spine by domain III, BMB Rep. 45 (2012) 227–232. doi: 10.5483/bmbrep.2012.45.4.227.

[47] F. C. J. Davies, et al., Endogenous epitope tagging of eEF1A2 in mice reveals early embryonic expression of eEF1A2 and subcellular compartmentalisation of neuronal eEF1A1 and eEF1A2, Mol Cell Neurosci. 126 (2023) 103879. doi: 10.1016/j.mcn.2023.103879.

[48] A. E. Perlegos, et al., Mettl3-dependent m(6)A modification attenuates the brain stress response in Drosophila, Nat Commun. 13 (2022) 5387. doi: 10.1038/s41467-022-33085-3.

[49] Y. E. Klingl, et al., Opportunities for histone deacetylase inhibition in amyotrophic lateral sclerosis, Br J Pharmacol. 178 (2021) 1353–1372. doi: 10.1111/bph.15217.

[50] O. R. Shanker, et al., Epigenetics of neurological diseases, Prog Mol Biol Transl Sci. 198 (2023) 165–184. doi: 10.1016/bs.pmbts.2023.01.006.

[51] T. Yang, et al., Epigenetic clocks in neurodegenerative diseases: a systematic review, J Neurol Neurosurg Psychiatry. 94 (2023) 1064–1070. doi: 10.1136/jnnp-2022-330931.

[52] L. L. Corey, et al., Localized recruitment of a chromatin-remodeling activity by an activator in vivo drives transcriptional elongation, Genes Dev. 17 (2003) 1392–1401. doi: 10.1101/gad.1071803.

[53] L. François-Moutal, et al., Heat shock protein Grp78/BiP/HspA5 binds directly to TDP-43 and mitigates toxicity associated with disease pathology, Sci Rep. 12 (2022) 8140. doi: 10.1038/s41598-022-12191-8.

[54] B. D. Freibaum, et al., Global analysis of TDP-43 interacting proteins reveals strong association with RNA splicing and translation machinery, J Proteome Res. 9 (2010) 1104–1120. doi: 10.1021/pr901076y.

[55] A. N. Coyne, et al., Post-transcriptional Inhibition of Hsc70-4/HSPA8 Expression Leads to Synaptic Vesicle Cycling Defects in Multiple Models of ALS, Cell Rep. 21 (2017) 110–125. doi: 10.1016/j.celrep.2017.09.028.

[56] C. Hubbert, et al., HDAC6 is a microtubule-associated deacetylase, Nature. 417 (2002) 455–458. doi: 10.1038/417455a.

[57] W. Tan, et al., Role of mitochondria in mutant SOD1 linked amyotrophic lateral sclerosis, Biochim Biophys Acta. 1842 (2014) 1295–1301. doi: 10.1016/j.bbadis.2014.02.009.

