## Supplementary Fig. 1, Fig. 2 and Table 1 for "Impact of histone deacetylase inhibition and arimoclomol on heat shock protein expression and disease biomarkers in primary culture models of familial ALS"

### Fernandez et al. Supplementary file

**Supplementary Figure 1.** Sequence of Stellaris probes to mus musculus *Hspa1a* and *Hspa8* mRNA labeled with Quasar 570 or Quasar 670 (Jacob-Tomas et al., 2022)

***Hspa1a* mRNA probes**
ttggttctgagtagctgtca, gctctgcttctcttgtcttc, tcgttggcgatgatctccac, gtcgaacacggtgttctgcg, ttgtagttcacctgcacctt, ggaagaacgaccggctctcg, accatggacgagatctcctc, agcgatctccttcatcttcg, gtgatcaccgcgttggtcac, cgctgagagtcgttgaagta, cacgtttagaccggcgatca, cgtgggctcgttgatgatcc, gtcgaagatgagcacgttgc, tcgatcgtcaggatggacac, tggccttcacctcgaagatg, tccacgaagtggctcaccag, ctggacgacagcgtcctctt, ctgagcacagctcttcgaac, atctgcgccttgtccatctt, ttgaagaagtcctgcagcag, ttgatgctcttgttcaggtc, tgcacgttctccgacttgtc, cttgatgagcgccgtcatca, ctcgtacacctggatcagca, ggatgccgctcagttcgaag, gaaggtcacctcgatctgtg, tgacgttcaggatgccgttg, ctgagcttgcccttgagacc, cagcaccttcttcttgtcag, gagatgacctcctggcactt,, cctggtacagcccactgatg, ctgagccagaggctcccttg, cagcagaggcctctaatcca atatctgtctcctagccagc, ataatgacagtcctcaaggc, acttgattgcaggacaaact, gataaagcccacgtgcaata, gacacacttgcagtgttaat, aaaacaaatcacatcagcgg, ggagagtccaaacacaaaga, atgtatcaacaatgtggccc, ctctgaaggacccgacacaa, agtctccgctgtcagtaatc, taaacgcaaggagaagcagc, gtgcaaccaccatgcaagat, caacctggaaacaagtccta, aaaactgaacacatgctggt, cggtaaaaattgacccgagt.

**Hspa8 mRNA probes**
Ttagacatggttgcttgtgt, gagatcaatgccaactgcag, ggcaataatttccacctttc, aaagcaacatagcttggcgt, cattgcaacctgattcttgg, gatcagacgtttggcatcaa, cagcatcatcaaacctacgc, atcattcaccaccatgaagg, tagaaactttttgtctcccc, agaaccatggaggacacttc, ctgcaatttccttcatcttt, atagttccagcatcttttg, tcgaagtacattgaggccag ctccgaccttcttatctaag, gacacatcaaaagtgccacc, aatcttctccacctaagtgg, atgattgaccattcggttgt, ctttcgcttgaactcagcaa, cttgttctcactgatgtctt, tcaatctcaatactggcctg, gtcaattccctcatagagag, cggaacaggtcagcattcaa, tcatggatctgtgacttgtc, gaatcttggggattctggta, cagctcttttccattgaaga, aacagcttcatcggggttaa, cagacttgtctccagataga, agcagcaaatcctgaacgtt, gggaaagaggagtgacatcc, gcttgatgaggacagtcatg, gtggtgaaagtctgtgtctg, tcaccttcatacacctgaat, gtgagctcgaactttccaag, gtaacctcaatctgagggac, gcagaaacattgaggatgcc, tggtgatggtgatcttgttc, gctcaatatcttccttactc, ttgtacttctcagcttcttg, gtgagttcttggaggaaacc, ttgctttcatgttgaaggca, ttgccttgaagtttctcatc, agaatcttctgtttgtcctc, gctgatgatttcattgcact, ctctggtacagcttggtaat, tgaagaagcaccaccagatg,
gacttaatccacctcttcaa, caaagctacaccttcttgga, tccatgttacttgttttggg

Jacob-Tomas, S., Alagar Boopathy, L.R., and Vera, M. (2022). Using Single-Molecule Fluorescence Microscopy to Uncover Neuronal Vulnerability to Protein Damage. Methods Mol Biol *2515*, 237-254.

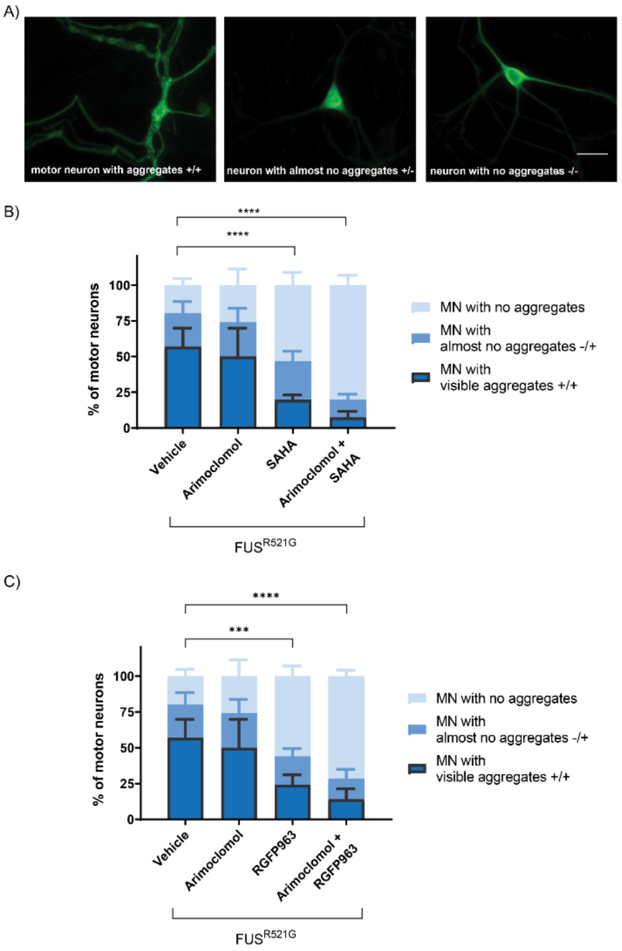

**Supplementary Figure 2.** The impact of HDAC inhibition, arimoclomol, and combination treatment on aggregate formation in motor neurons expressing $FUS^{R521}$. $FUS^{R521G}$ was expressed in motor neurons of dissociated spinal cord-DRG cultures by intranuclear microinjection of plasmid vector. Cultures were treated with vehicle (DMSO), 4 μM arimoclomol, or HDAC inhibitor (4μM SAHA or 1μM RGFP963) alone or in combination with arimoclomol for three days. Cultures were fixed and immunolabeled with FUS antibody. **A** The proportion of motor neurons exhibiting cytoplasmic aggregates was quantified as shown the micrographs: neurons with multiple aggregates throughout the cytoplasm (+/+), neurons with few aggregates (*/-), or neurons without visible aggregates (-/-). **B** SAHA and **C** RGFP963, alone and in combination with arimoclomol, but not arimoclomol alone, reduced the percentage of neurons with cytoplasmic aggregates. Data are presented as mean ± S.D., n = 6-7 cultures per group. Statistical significance was evaluated through one-way ANOVA followed by Bonferroni post hoc analysis. ***$p<0.001$ and ****$p<0.0001$. Scale bar = 20μm.

**Supplementary Table 1.** Summary of findings. → no significant change; ↑increase; ↓decrease; n.d. not determined. Included are findings from this study, Tradewell et al., 2011 [24], Tibshirani et al., 2015, 1017 [35,13] and Kuta et al., 2020 [7].

| Parameter | Treatment | ALS Variant | | |
|---|---|---|---|---|
| | | TDP-43$^{G348C}$ | FUS$^{R521G}$ | SOD1$^{G93A}$ |
| HSPA1A | Vehicle | ↑ | → | ↑ |
| | SAHA | ↑ | → | ↑ |
| | RGFP109 | ↑ | → | ↑ |
| | RGFP963 | ↑ | → | |
| | Tubastatin A | → | → | → |
| | Arimoclomol | → | → | ↑ |
| | Arimoclomol + HDAC inhibitor (SAHA, RGFP963 or RGFP109 | ↑ | → | ↑ |
| *Hspa1a* mRNA | Vehicle | → | → | → |
| | RGFP963 | → | → | ↑ |
| | Arimoclomol | → | → | ↑ |
| | RGFP963 + Arimoclomol | → | → | → |
| *Hspa8* mRNA soma | Vehicle | ↓ | ↓ | ↓ |
| | RGFP963 | → | → | ↑ |
| | Arimoclomol | → | → | → |
| | RGFP963 + Arimoclomol | ↑ | → | ↑ |
| *Hspa8* mRNA dendrite | Vehicle | ↓ | ↓ | ↓ |
| | RGFP963 | → | → | ↑ |
| | Arimoclomol | → | → | → |
| | RGFP963 + Arimoclomol | → | → | ↑ |
| Histone acetylation | Vehicle | ↓ | ↓ | n.d. |
| | SAHA | ↑ | ↑ | |
| | RGFP963 | ↑ | | |
| | Arimoclomol | → | → | |
| | SAHA + Arimoclomol | ↑ | | |
| Nuclear TDP-43 or FUS | Vehicle | ↓ | ↓ | n.d. |
| | SAHA | ↑ | ↑ | |
| | RGFP109 | ↑ | ↑ | |
| | RGFP963 | ↑ | | |
| | Tubastatin A | → | → | |
| | Arimoclomol | ↑ | ↑ | |
| | Arimoclomol + (SAHA, RGFP 109 or RGFP963 | ↑ | ↑ | |
| Nuclear Brg1 | Vehicle | ↓ | ↓ | n.d. |
| | SAHA | ↑ | | |
| | RGFP963 | ↑ | | |
| | Arimoclomol | ↑ | | |
| | RGFP963 + Arimoclomol | ↑ | | |
| Mitochondrial transport | Vehicle | ↓ | ↓ | ↓ |
| | RGFP963 | → | | → |
| | Arimoclomol | → | | → |
| | RGFP963 + Arimoclomol | → | → | ↑ |
